# Oncomimetic β-catenin activity onset, duration and region defines aberrant intestinal development

**DOI:** 10.64898/2026.05.07.723518

**Authors:** Birga Soetje, Hui Ma, Sarah Imtiaz, Agustin Corbat, Hernan E Grecco, Lisaweta Schröder, Yannick Brüggemann, Sabrina Seidler, Michael Reichl, Philippe IH Bastiaens

## Abstract

In early intestinal carcinogenesis, adenoma formation is commonly initiated by loss-of-function mutations in a tumor suppressor that lead to oncoprotein gain-of-function, like in the tumor suppressor–oncoprotein pair APC–β-catenin. Small intestinal organoids provide an *in vitro* system to study consequences of such mutations on tissue organization. However, conventional genetic manipulations do not allow precise control over the onset and duration of oncoprotein activity to study their influence on tissue transformation. Furthermore, homogenous tissues of clonal genetic models do not readily capture cellular interaction among mutated and neighboring wildtype tissue during early transformation. To mimic oncogenic activation of β-catenin, we established a chemical-(opto)genetic approach to gain bio-orthogonal acute, spatial and temporal control over β-catenin oncoprotein stability and relate oncoprotein levels to morphological development of mouse small intestinal organoids.

We identified aberrant phenotypes that result from bio-orthogonally induced oncoprotein activity but persist even after oncoprotein depletion. Furthermore, local activation of oncomimetic β-catenin activity within the stem cell niche leads to aberrant differentiation during homeorhesis and homeostasis, recapitulating early events of tissue transformation.

## Introduction

In the well characterized progression of sporadic colorectal cancer (CRC), the accumulation of loss and gain of function mutations causes carcinogenesis following the conventional adenoma to carcinoma sequence (Fearon & Vogelstein, 1990; Leslie *et al*, 2002). The frequently occurring initiating loss-of-function of the tumor suppressor APC (adenomatous polyposis coli) causes stabilization of β-catenin leading to crypt dysplasia and early adenoma formation (Andreu *et al*, 2005; Fearon, 2011; Harada *et al*, 1999). These morphological transformations arise from the deregulation of two highly characterized orthogonal functions of β-catenin : 1) as transcriptional cofactor for T-cell factor/Lymphoid enhancer-binding factor (TCF/Lef) (Behrens *et al*, 1996; Huber *et al*, 1997; Molenaar *et al*, 1996) regulating cell proliferation (Hatzis *et al*, 2008) and 2) in cell-cell adhesion where β-catenin is part of the adherens junction complex (Ozawa *et al*, 1989) affecting tissue morphology and epithelial-to-mesenchymal transition in late stages of carcinogenesis (Kim *et al*, 2019; Valenta *et al*, 2012; van der Wal & van Amerongen, 2020). Wnt-binding to the receptor complex of frizzled proteins and LRP5/6 enables the scaffold protein APC in the destruction complex to control cytoplasmatic β-catenin levels (Clevers & Nusse, 2012; Li *et al*, 2012).

The oncogenic β-catenin gain-of-function can arise on the one hand upon loss-of-function mutations of APC (Li *et al*., 2012) impairing the continuous synthesis/degradation cycle. On the other hand hotspot mutations in β-catenin exon3 (Kim & Jeong, 2019) transform β-catenin from a controllable proto-oncoprotein to an oncoprotein by impairing phosphorylation by members of the destruction complex CKIα (casein kinase 1α; S45) and GSK3β (glycogen synthase kinase 3β; T41/S37/S33) (Amit *et al*, 2002; Huels *et al*, 2015; Liu *et al*, 2002) and subsequent ubiquitination or neddylation and proteasomal degradation of β-catenin by β-TrCP1/2 E3 ubiquitin-ligases (MacDonald *et al*, 2009; Wang *et al*, 2022).

Intestinal organoids have emerged as powerful *in vitro* systems to study epithelial organization and regeneration and that have provided a wealth of knowledge on intestinal tumorigenesis in relation to genomic mutations such as loss of APC (Clevers, 2026; Huels *et al*., 2015; Sato *et al*, 2009). During organoid culture, seeded intestinal stem cells (ISC) and crypts reproducibly recapitulate a process of repair after injury and cellular differentiation by which small intestinal organoids (re-)develop a prototypical compartmentalized intestinal crypt-villus structure (Sato & Clevers, 2013; Sato *et al*., 2009), resembling homeorhesis (Ray, 2021). Recapitulating the native tissue, crypt regions harbor the ISC and Paneth cells (PC) at the base and a zone of transient amplifying cells that continuously migrate upwards along the crypt-villus axis and differentiate into enterocytes and secretory cells in the villus region (Sato & Clevers, 2013; Serra *et al*, 2019). Intestinal organoids thereby have provided a model system to study consequences of oncogenic mutations on cell differentiation and tissue organization (Date & Sato, 2015; Drost & Clevers, 2018; Merker *et al*, 2016). However, during organoid culture the stem cells propagate oncogenic mutations to all cells of the developing and differentiating organoid leading to clonal expansion over multiple rounds of propagation in culture. These genetically homogenous tissues do therefore not recapitulate initial events of tumorigenesis within a healthy tissue in which biochemical and biomechanical cell-cell communication affects the development of the tumor. Furthermore, constitutive oncoprotein expression does not enable the assessment of the changes that would occur after loss of the initiating oncoprotein.

To overcome these limitations of conventional genetic approaches, we developed a chemical genetic approach in mouse small intestinal organoids (mSIOs) that not only enables global, acute and reversible chemical control but also local, opto-chemical control of β-catenin oncoprotein activity. This system enables precise control over the onset, duration, and spatial confinement of oncogenic β-catenin activity and thereby allows to address the initiation and intercellular communication among transformed and healthy cells during tissue transformation in an organoid. With this bio-orthogonal, opto-chemical approach we addressed the following questions: 1) Transformational ability: is oncogenic β-catenin activity sufficient to induce aberrant tissue development and how does the developmental stage at activation influence this outcome. 2) Reversibility: is development of an aberrant morphological phenotype dependent on sustained oncoprotein activity? and 3) Propagation: can local oncoprotein activity within cellular niches of developing crypts transform part or the whole of the normal organoid to aberrant tissue?

To answer these questions, we identified and mapped normal and aberrant organoid phenotypes to a morphometric readout which allows to classify organoid states and recapitulate their developmental trajectories. We found that globally induced oncoprotein activity causes an aberrant developmental trajectory distinct from that of constitutive oncoprotein expression and this acutely induced aberrant trajectory is not dependent on constant presence of the initiating oncoprotein. Modulation of onset showed that these aberrant trajectories are dependent on the organoid organization at activation. However, the common resulting feature during late onset of oncoprotein activity is an increase in crypt-like precursor regions either by augmented crypt size or number. Local opto-chemical oncoprotein activation in developing crypt precursor regions during organoid regeneration had the same state-at-onset dependence as global, chemical oncoprotein stabilization with the main outcome of elevated crypt fission. In contrast, oncoprotein activation in mature crypts during organoid homeostasis led to elevated crypt proliferation and impaired cell migration and differentiation, resembling the effects caused by loss of APC. Importantly, the local activation also induced a destabilization of the organoid tissue, causing crypt separation from the originating tissue.

In the tissue context, these results indicate a capability of oncogenic β-catenin to induce dysplasia-like effects not only in mutated crypts but also by long-range impairment in neighboring tissue regions in conditions of tissue repair after injury.

These findings establish our opto-genetic system of oncomimetic β-catenin activity as an effective and tunable model to study tumorigenesis.

## Results

### Genetically encoded opto-chemical β-catenin oncoprotein activity control

To establish measurable bio-orthogonal (opto)-chemical control over the stability and therefore oncogenic β-catenin activity in space and time, a polycistronic PiggyBac plasmid was designed that encodes _(I)_ a fluorescent (mCitrine) β-catenin4A-mCitrine-DD fusion protein (βCtn4A-DD, 1184 aa, Mw: 131.4 kDa), _(II)_ a photoactivatable mCherry (PAmCherry) and _(III)_ a nuclear targeting Halo-3xNLS for compound retention (Fig. 1A). β-catenin4A contains alanine mutations at four CKIα/GSK3β regulatory phosphorylation sites (S33A/S37A/T41A/S45A) of β-catenin (Amit *et al*., 2002; Liu *et al*., 2002; Wu & He, 2006), bypassing the Wnt-controlled APC destruction complex-mediated degradation of β-catenin. β-catenin4A fusion to ecDHFR^N18T/A19V^ destabilization domain (DD; *E. coli* dihydrofolate reductase) (Iwamoto *et al*, 2010) causes constitutive degradation of the fusion protein, enabling bio-orthogonal control of βCtn4A-DD stability by binding of the small molecule trimetoprim (TMP) to DD (Fig. 1B). This system enables two modes of control of βCtn4A-DD activity: _(i)_global, reversible stabilization by addition and withdrawal of TMP; _(ii)_acute, local stabilization by 405 nm irradiation of photo-caged TMP (NvocTMP-Cl) (Chen *et al*, 2017) in selected cells. Compound retention of NvocTMP-Cl within individual cells is mediated by covalent binding of the chlorohexyl (Cl) moiety to co-expressed Halo-3xNLS (Fig. 1C) which hinders diffusion of caged and de-caged dimerizer, ensures nuclear targeting of the oncoprotein. Co-expression of the three contruct components is facilitated via ribosomal skipping sites (P2A-T2A) (Donnelly *et al*, 2001a; Donnelly *et al*, 2001b; Liu *et al*, 2017; Pan *et al*, 2017). The co-expressed photoactivatable mCherry (PAmCherry) (Subach *et al*, 2009) enables immediate tracking of cells in which NvocTMP-Cl was uncaged by 405 nm light. The expression plasmid is termed bCtCDD-PCh-Hal and further designated as bCtC-DD (Fig.1A).

**Figure 1.**
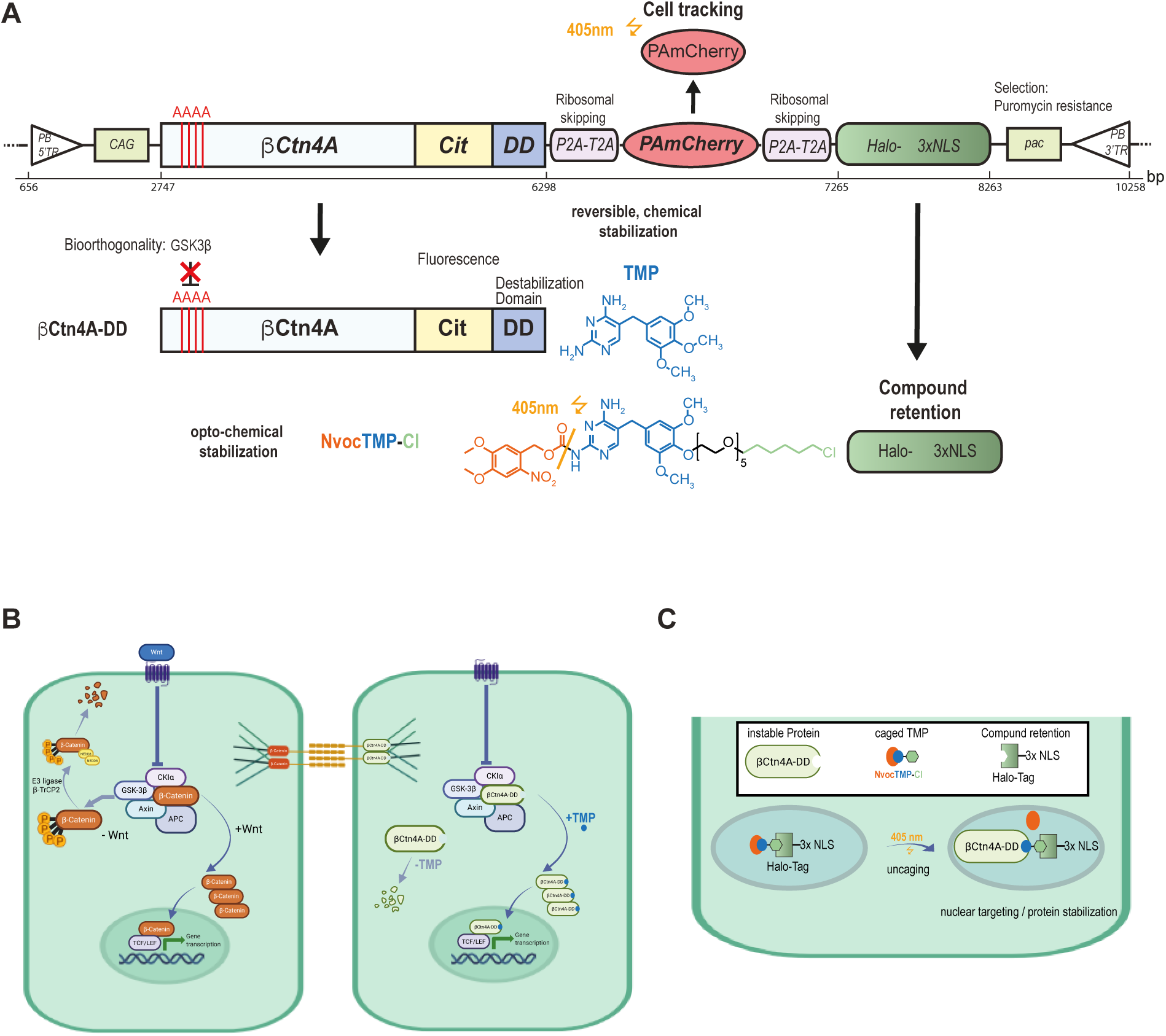
Design of opto-chemical control of oncogenic β-catenin. **A.** Polycistronic PiggyBac construct (*bCtC-DD*) encoding fluorescent βCtn4A-DD, PAmCherry for Cell Tracking and Halo-3xNLS for NvocTMP-Cl retention. bp: base pairs. From left to right: *PB 5’TR*: 5’ PiggyBac inverted terminal repeat; *CAG*: human cytomegalovirus early enhancer/chicken beta actin promotor; *bCtn4A*: β-catenin sequence with S33A/S37A/T41A/S45A mutations; *Cit*: monomeric yellow fluorescent protein Citrine sequence; *DD*: ecDHFR N18T/A19V destabilization domain sequence based on E.coli dihydrofolate reductase; *P2A-T2A*: first ribosomal skipping site; *PAmCherry*: photoactivatable monomeric red fluorescent protein Cherry; *P2A-T2A*: second ribosomal skipping site; *Halo-3xNLS*: sequence encoding haloalkane dehalogenase fused to 3x nuclear localization signal; *pac*: Internal ribosomal entry site and puromycin N-acetyltransferase expression cassette for selection; *PB 3’TR*: 3’ PiggyBac inverted terminal repeat. Black arrows: transcription/translation. TMP: trimethoprim for reversible chemical stabilization. NvocTMP-Cl: 6-nitroveratryloxycarbonyl (Nvoc) caged TMP conjugated to chlorohexyl moiety (Cl) for opto-chemical stabilization by 405 nm light irradiation. Cl-moeity covalently binds to Halo-3xNLS for compound retention. **B.** Left cell: Wnt-dependent regulation of β-Catenin destruction cycle. Without Wnt, β-Catenin is bound by the destruction complex, phosphorylated by GSK-3β and CK1α and further degraded by E3 ligases as β-TrCP2. Upon Wnt binding, Axin is recruited to the Frizzled-LRP5/6 complex, disassembling the destruction complex, which enables cytoplasmic accumulation of β-Catenin and nuclear translocation to form TCF/LEF- β-Catenin transcription complex. Additionally, β-Catenin is found in adherens junctions. Right cell: βCtn4A-DD is instable in absence of TMP. Addition of TMP leads to stabilization and subsequent accumulation and nuclear translocation of βCtn4A-DD. βCtn4A-DD can still be bound by the destruction complex and adhesion junctions. **C** Schematic representation of the principle for opto-chemical protein stabilization. The Halo-3xNLS fusion protein is retained in the nucleus of expressing cells. The caged compound NvocTMP-Cl covalently binds to Halo-3xNLS and is thus retained in the nucleus of the cell. 405 nm irradiation leads to photo-uncaging of the Nvoc group enabling the interaction and of the nuclear TMP-Cl with the ecDHFR-based destabilization domain (DD) and thereby stabilization of the oncoprotein βCtn4A-DD. Local oncoprotein stabilization is enforcing nuclear βCtn4A-DD accumulation.

To validate chemical control over βCtn4A-DD stabilization/destabilization, βCtn4A-DD levels were compared to endogenous β-catenin (βCtn) in Hela cells stably expressing βCtn4A-DD. Quantitative western blot analysis showed that after 48 h TMP (10 µM) incubation (TMP[48]), βCtn4A-DD increased ∼1.8±0.6-fold (mean±SD) with respect to basal levels, whereas endogenous βCtn remained unchanged (Fig. 2A, EV2A). In contrast, inhibition of the APC destruction cycle with the GSK3β inhibitor CHIR99021 (6 µM) led to a ∼3.9±0.9-fold (mean±SD) increase in endogenous βCtn but did not affect βCtn4A-DD (Fig. 2A, EV2A), confirming bio-orthogonal chemical control of βCtn4A-DD levels by TMP. βCtn4A-DD also exhibited intact transcriptional activity as apparent from the comparable ∼5.5±1.8-fold (mean±SD) increase in Axin2 levels after TMP[48] or βCtn stabilization by CHIR99021 (Fig. 2B, EV2B). Live cell imaging showed that βCtn4A-DD fluorescence gradually increased within ∼10 h during TMP[48], whereas washout of TMP after 24 h (TMP[24,-24]) caused a decline of βCtn4A-DD fluorescence to the baseline of vehicle-treated (DMSO) cells within ∼4 h (Fig 2C,D, EV2C). The corresponding protein level decline after TMP washout was also apparent from western blot analysis (Fig. 2A). These experiments in Hela cells thus confirm reversible bio-orthogonal chemical control of βCtn4A-DD oncoprotein stability.

**Figure 2.**
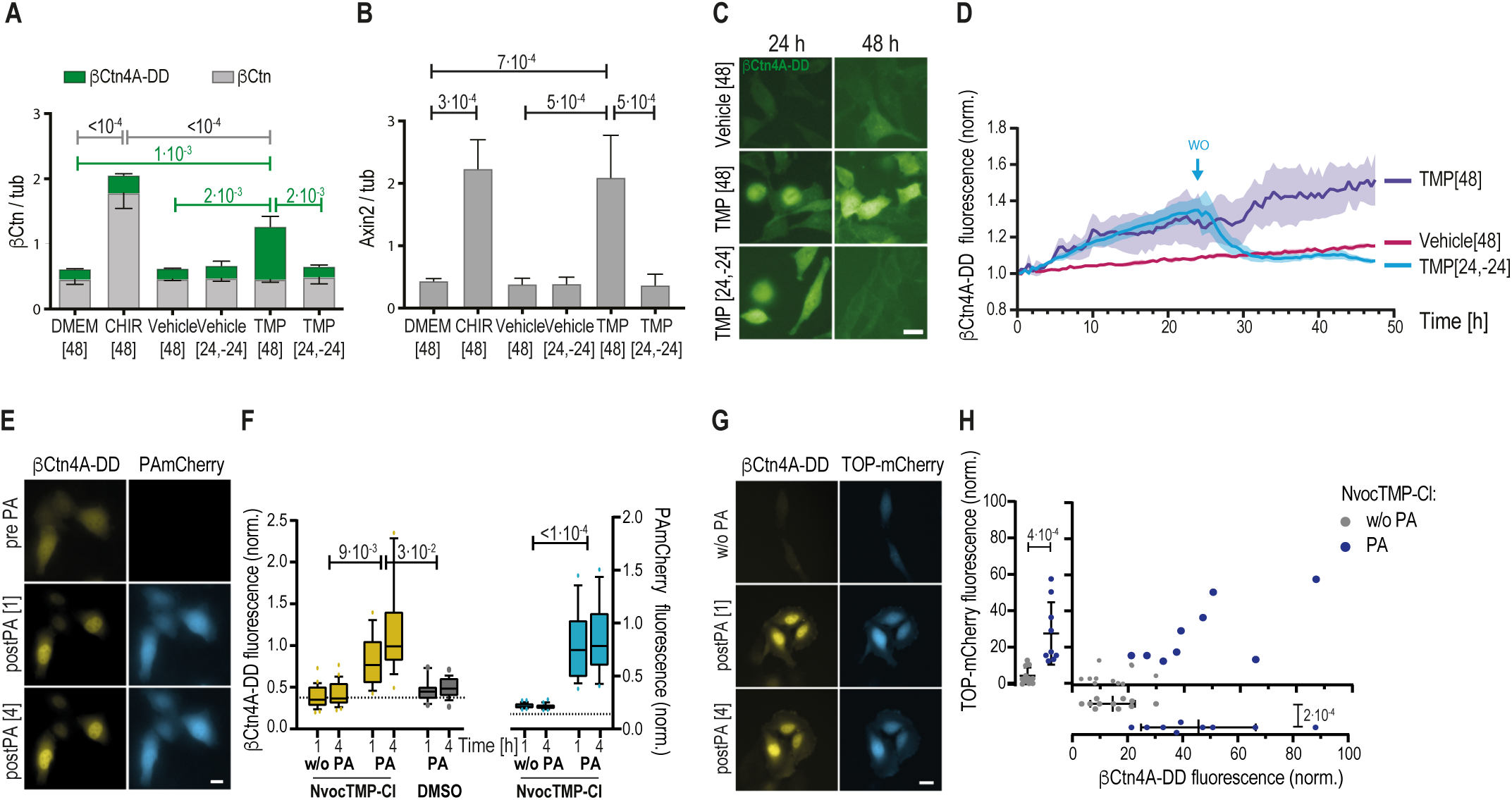
Bioorthogonal opto-chemical oncogenic β-Catenin control in HeLa cells. **A.** Relative protein levels (βCtn/tub) in transgenic HeLa bCtC-DD normalized to Tubulin as determined by western blot. βCtn4A-DD/Tubulin (green), βCtn/Tubulin (grey). DMEM[48]: 48 h full growth media; CHIR[48]: 48 h DMEM + 5 µM CHIR99021; Vehicle[48]: 48 h DMEM + 0.1% DMSO; Vehicle[24,-24]: 24 h DMEM + 0.1% DMSO followed by washout and 24 h DMEM; TMP[48]: 48 h DMEM + 10 µM TMP; TMP[24,-24]: 24 h DMEM + 10 µM TMP followed by washout and 24 h growth in DMEM. Mean±SD, N=3. p-values (color coded): two-way ANOVA, Tukey’s multiple comparison test. **B.** Corresponding relative Axin2 levels normalized to Tubulin (Axin2/tub). Mean±SD, N=3. p-values: one-way ANOVA, Tukey’s multiple comparison test. **C.** Representative micrographs of βCtn4A-DD citrine fluorescence at indicated time points in HeLa bCtC-DD. Vehicle[48], TMP[48], TMP[24,-24] as described in (A). scale bar: 20 µm. **D**. Corresponding normalized mean βCtn4A-DD citrine fluorescence ± SEM over time (N=2, n=4/condition). **E**. Representative micrographs of βCtn4A-DD and PAmCherry in HeLa bCtC-DD after NvocTMP-Cl incubation for 60 min (prePA), 1 h (postPA[1]) and 4 h (postPA[4]) after uncaging by 405 nm irradiation for 40 s (6 J/cm^-2^). **F**. Quantification of βCtn4A-DD (left) and PAmCherry (right) as normalized average fluorescence/cell. NvocTMP-Cl: Cells preincubated with NvocTMP-Cl (10 μM); DMSO: cells preincubated with 0.1% DMSO vehicle. Boxplot: 10-90% with outliers. Dotted line: mean intensity prePA. n=17-33 cells. p-values: One-way ANOVA, Tukey’s multiple comparisons test. **G**. Representative micrographs of βCtn4A-DD and TOP-mCherry transcriptional reporter fluorescence in HeLa bCtC-DD-Hal, lacking PAmCherry, after NvocTMP-Cl incubation for 60 min, without decaging (w/o PA), 1 h (postPA[1]) and 4 h (postPA[4]) after 405 nm irradiation for 40 s (6 J/cm^-2^). **H**. Corresponding scatterplots of normalized mean βCtn4A-DD and TOP-mCherry fluorescence per cell. n= 9-11 cells, 4 timepoints/cell (1, 2, 3, 4h). Blue dots: PA, Grey dots: w/o PA. Experimental variance parallel to axes: Mean±SD, p-values: One-way ANOVA.

Opto-chemical control of βCtn4A-DD stabilization in transgenic HeLa bCtC-DD was assessed by 1 h preincubation with NvocTMP-Cl (10 µM), washout, and irradiation with 405 nm light (6 J/cm²). Nuclear βCtn4A-DD stabilization was detectable as early as 1 hour after irradiation and increased further until ∼4 h relative to the low basal levels in non-irradiated cells or irradiated vehicle-pretreated controls (Fig. 2E,F). PAmCherry fluorescence exhibited a similar increase at 1 h and 4 h after 405 nm light exposure in relation to the pre-activation signal, establishing its functionality as maker for irradiated and thereby activated cells (Fig. 2E,F). In Hela cells that lacked the PAmCherry coding sequence but co-expressed TOP-mCherry, a reporter for TCF/Lef promotor activation (Veeman *et al*, 2003), opto-chemically stabilized βCtn4A-DD fluorescence levels correlated with TOP-mCherry levels, indicating functional transcriptional activity (Fig. 2G,H).

This shows that the activity of the (opto-)chemically stabilized oncoprotein βCtn4A-DD effectively mimics acute oncogenic mutations and can thereby be characterized as oncomimetic.

### Morphometric classification of aberrant mSIO development

We proceeded to test the bio-orthogonal, chemical control of βCtn4A-DD stabilization in transgenic small intestinal organoids (mSIOs) incorporating the bCtC-DD construct. Quantitative Western blot analysis showed that TMP-treatment for 144 h (10 µM; TMP[144]), specifically increased βCtn4A-DD levels to 4.3±2.3 (mean±SD) that of endogenous βCtn, which showed a 2-fold reduction (0.5±0.4; mean±SD) compared to Vehicle[144]. In contrast, Wnt3a-treatment with Wnt3a-conditioned media for 144 h (Wnt[144]; 50% Wnt3a-CM) specifically increased endogenous βCtn to 2.05±0.37 (mean±SD) times that of basal βCtn protein levels without affecting βCtn4A-DD levels (Fig 3A; Fig EV3A). TMP-stabilized βCtn4A-DD levels correlated with TCF/Lef-transcriptional activity measured using a TOP-mCherry transcriptional reporter (Veeman *et al*., 2003) and demonstrated the chemically inducible transcriptional activity of βCtn4A-DD in mSIOs (Fig 3B, EV3B). Together these results confirm the bio-orthogonal control of βCtn4A-DD stabilization, transcriptional activity and functional negative feedback affecting endogenous βCtn levels.

**Figure 3.**
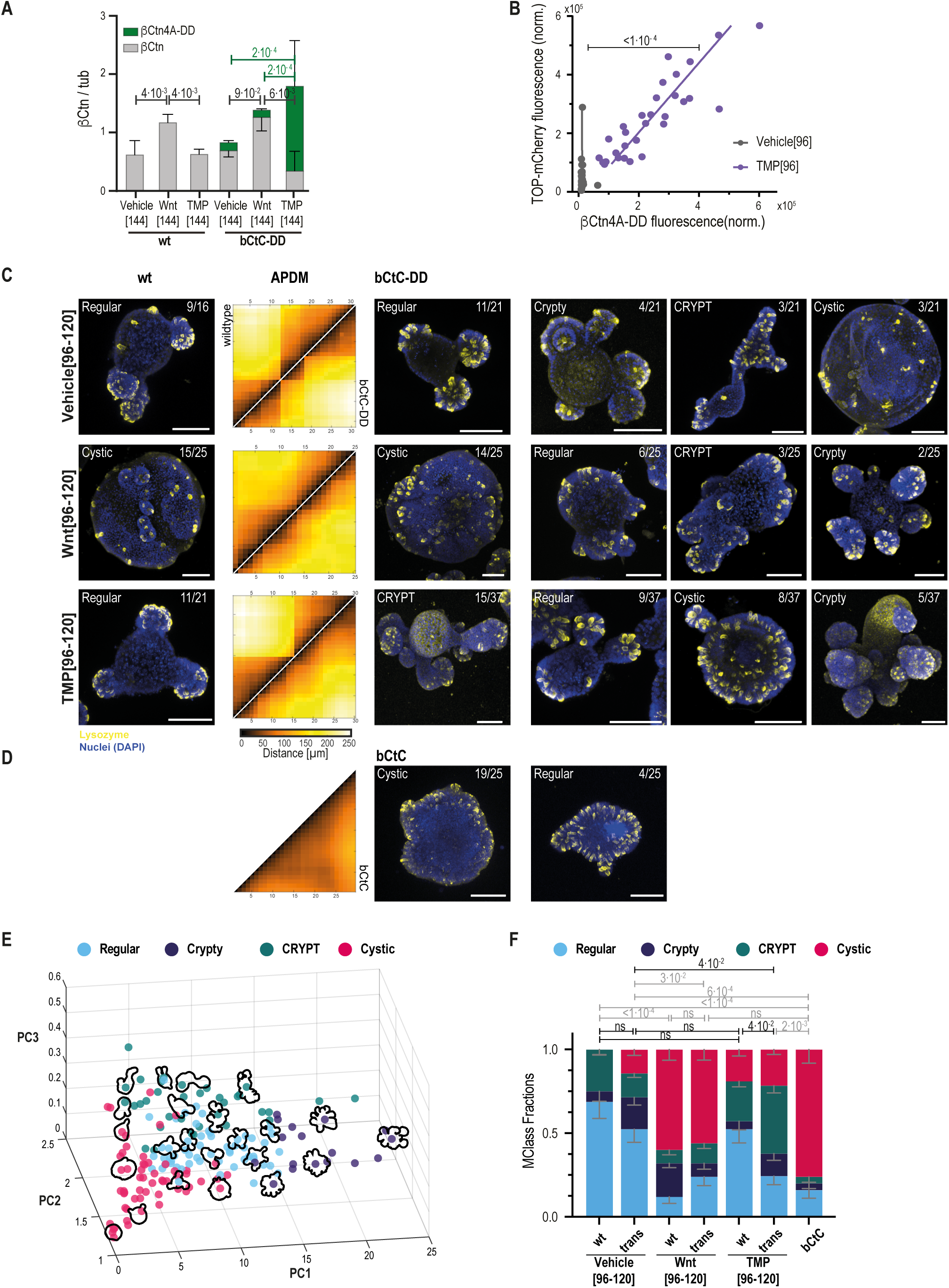
Morphometric classification of normal versus aberrant mouse small intestinal organoids (mSIOs). **A.** Relative protein levels (βCtn/tub) of βCtn4A-DD /Tubulin (green) and endogenous β-Catenin βCtn/Tubulin in wildtype (wt) and transgenic bCtC-DD mSIOs from western blots. Vehicle[144]: 144 h ENR + 0.1% DMSO, Wnt[144]: 144 h ENR + 50% Wnt3a-conditioned media, TMP[144]: 144 h ENR + 10 µM TMP. Mean±SD, N=4, p-values (color coded): wt: one-way ANOVA; bCtC-DD: two-way ANOVA; Tukey’s multiple comparison test. **B.** Scatterplot of βCtn4A-DD versus TOP-mCherry fluorescence intensity from crypt-like structures of bCtC-DD mSIOs grown in Vehicle[96] (grey dots) or TMP[96] (purple dots). Corresponding colored lines: linear fit to data; TMP[96] Spearman’s r= 0.8776, vehicle[96] Spearman’s r= 0.3095 p: ANCOVA. Vehicle[96]: N=2, n=23; TMP[96]: N=2, n=14. **C.** Representative maximum intensity projections (MIPs) of 3D CLSM immunofluorescence stacks of mSIOs and corresponding APDMs. Yellow: Paneth cell marker antibody α-lysozyme, Blue: Nucleus marker DAPI. Left Column (wt): Wildtype mSIOs; Right block (bCtC-DD): bCtC-DD mSIOs. Rows from top to bottom: Vehicle[96-120]: 96-120 h ENR + 0.1% DMSO; Wnt[96-120]: 96-120 h ENR + 50% Wnt3a-conditioned media; TMP[96-120]: 96-120 h ENR + 10 µM TMP. APDM: corresponding average phase-matched distance matrices. Above diagonal: wt, N=4/3/4; n=16(2/4/2/8)/25(6/3/16)/21(1/1/2/17), Below diagonal: bCtC-DD, N=4/6/6; n=21(4/5/2/10)/25(3/3/4/1/2/12)/37(10/3/3/6/3/3/12). Color bar: PC distance (µm). Mclasses and corresponding fractions indicated at top of MIPs in white. Scale bar: 100 µm. **D.** Same as (C) for bCtC[96] mSIOs: 96 h ENR; N=3; n=25(3/9/13). **E.** 3D Principal component projection (PC1, PC2, PC3) of mSIO’s 8-dimensional morphometric parameter distribution (Fig. EV3H,I) with the identified Mclasses ‘Regular’ (Cyan), ‘CRYPT’ (green), ‘Crypty’ (violet), ‘Cystic’ (red). MIP contours of mSIOs within the morphometric space projection are shown. **F.** Mclass distributions of wildtype (wt) and bCtC-DD (trans) in Vehicle[96-120], Wnt[96-120], TMP[96-120], and bCtC[96] mSIOs. Mean±SD by Mclassifier confusion table (Fig. EV3J). N=3-6; n=16/21/25/25/21/37/25 (left to right; details in (C,D)). p-values: chi-squared test. ns: non significant.

To identify aberrant development of transgenic mSIOs that express a stable (bCtC; βCtn4A) or destabilized (bCtC-DD; βCtn4A-DD) version of oncomimetic active β-catenin, overall Paneth Cell (PC) organization was assessed. As direct neighbors of intestinal stem cells in the crypt base, PC organization is an indicator of overall organoid organization. To map the organoids PC organization, bCtC-DD, bCtC and control (wt) mSIOs were fixed after 96-120 h of development and PCs were stained by indirect immunofluorescence against the PC marker lysozyme (Clevers & Bevins, 2013) and inter-PC distances were obtained from 3D confocal micrographs (Fig. 3C, Fig. EV3C). Proximity sorted PC distances result in one Distance Matrix (DM) for each mSIO (Fig. EV3D; methods) and represent single organoid organization. A binarized version of the DMs is used to assess cluster regularity based on the Shannon Entropy (SE) of binary DMs (Fig. EV3E; methods). In addition, Average Phase-matched Distance Matrices (APDMs; methods) represent the common PC-organization within a mSIO population (Fig. 3C,D) and allow the comparison of organoid populations.

The majority of bCtC-DD transgenic organoids developed a prototypical crypt-villus structure. Their APDM exhibited a pattern comparable to that of wt mSIOs showing well-separated PC clusters. Their SE distributions were statistically indistinguishable indicating PC organization in defined clusters of the crypt regions (Fig. 3C, Fig. EV3E).

Continuous exposure of transgenic and wt mSIOs to exogenous Wnt for up to 120 h (Wnt[96-120]) predominantly induced a cystic morphology (Heinz *et al*, 2020; Sato *et al*, 2011a) with comparable APDM patterns and SE distributions that are indicative of smaller, evenly distributed PC clusters as compared to controls.

This Wnt-induced aberrant development was in contrast to that of bCtC-DD mSIOs subjected to continuous TMP exposure for up to120 h (TMP[96-120]) in which was βCtn4A-DD stabilized. The latter predominantly displayed expanded crypts, resulting in a short-distance “pearls-on-a-string” APDM pattern and a significantly increased SE distribution which is distinct with respect to controls (Fig. 3C, Fig. EV3E).

Although SE distributions were similar between TMP- and Wnt-treated mSIOs, this is not reflected by the morphology. In particular, APDMs suggest increased PC cluster sizes upon TMP exposure compared to Wnt. These TMP-induced APDM pattern and SE distributions in bCtC-DD mSIOs, however, clearly indicate enhanced PC cluster irregularity as compared to wt mSIOs exposed to TMP. The latter had a PC-organization similar to wt mSIOs, showing that TMP (10 μM) exposure by itself does not cause aberrant development. Furthermore, bCtC mSIOs that stably express βCtn4A oncoprotein, completely lost the crypt-villus structure with no apparent pattern in the APDM (Fig. 3D) and the most elevated SE distribution (Fig. EV3E). Thus, early induction of oncomimetic βCtn4A-DD activity in the entire developing organoid led to aberrant PC cluster arrangement that is distinct from the complete loss of PC-organization resulting from clonal expansion of chronic oncomimetic βCtn4A expression.

To establish a classification of normal and aberrant mSIO morphologies, treatment-related central Paneth cell organizations within the mSIOs were identified (Fig. EV3 F,G; methods) and mapped into an optimized 8-dimensional morphometric space that is invariant to scale, rotation and translation (Mspace; Fig. EV3H,I; methods). This Mspace is visualized using a principal components projection (PC1, PC2, PC3). Within this Mspace, the distributions of bCtC and controls (wt Vehicle, wt TMP, bCtC-DD Vehicle) served as benchmark classes of the aberrant ‘Cystic’ and prototypical ‘Regular’ mSIO morphologies, respectively. TMP and Wnt exposure in wt and bCtC-DD mSIOs led to three additional morphological clusters which are characterized by enlarged, enriched, or absence of crypts (Fig. EV3I; methods). To now classify aberrant versus normal morphologies, a decision-tree based morphological classifier (Mclass) was generated by supervised training with the identified clusters as ground truth (Fig. EV3I). In this iterative process it turned out that the fusion of the ‘bCtC’ and the ‘absence of crypts’ clusters increased overall classifier accuracy. The resulting classifier with the four distinct morphological classes (Mclasses) of ‘Regular’, ‘CRYPT’, ‘Cystic’ and ‘Crypty’ converged to a mean accuracy of 0.94±0.02 and an F1-Score of 0.87±0.02 (Fig. EV3J).

In the principal components projection (PC1, PC2, PC3) the mSIOs are colored by their four Mclasses (Fig. 3E) and shown together with exemplary organoid contours within each Mclass. From this projection it is apparent that the ‘Regular’ Mclass is characterized by the prototypical crypt-villus structures and is surrounded by three aberrant classes of which the ‘Cystic’ is featureless. The ‘CRYPT’ Mclass has an extended morphology with large crypt-like structures whereas the ‘Crypty’ Mclass has a more circularly arranged morphology with increased numbers of crypt-like structures around the organoid.

Mclass fraction distributions for each organoid population of a certain condition show that fully developed bCtC-DD transgenic mSIOs are statistically indistinguishable from wt and mostly manifested as ‘Regular’ (Fig. 3F). As expected (Heinz *et al*., 2020), Wnt[96-120] exposure predominantly induced ‘Cystic’ development with a minor population of ‘Crypty’. This ‘Crypty’ Mclass also occurred in the transgenic mSIOs but rarely in the wt controls. However, the difference in morphological outcomes of stable βCtn4A expression (‘Cystic’) and acute βCtn4A-DD stabilization by TMP (‘CRYPT’) indicated that the time of onset and duration of oncoprotein activity affects the trajectory of aberrant development (Fig. 3F). This question could now be investigated by relating βCtn4A-DD fluorescence to morphodynamic developmental trajectories during individual mSIO development .

### Oncomimetic β-catenin temporal profiles affect morphodynamic trajectories

The development of individual mSIOs within populations starting from disintegrated organoids (Fig. EV4A) was tracked by automated widefield transmission microscopy with a time interval of 6 h for up to 120 h. Contour development for each organoid was obtained by segmentation of the z-stack median projection and growth curves were determined as the median fold-change in cross-sectional organoid area relative to that at 24 h (Fig. EV4B). mSIO morphodynamics were then assessed by determination and clustering organoid contour trajectories according to their end-stage (96 h) classification in Mspace (Fig. EV4C; methods).

Principal component projections (PC1, PC2) of the trajectories (Fig. 4A-C; bottom) were generated together with corresponding Mclass fractions (Fig. EV4D). In addition, the developing contours of a single organoid most proximal to the main trajectory was determined to assess the morphodynamic transitions along that developmental path (Fig. 4A-C; top).

**Figure 4.**
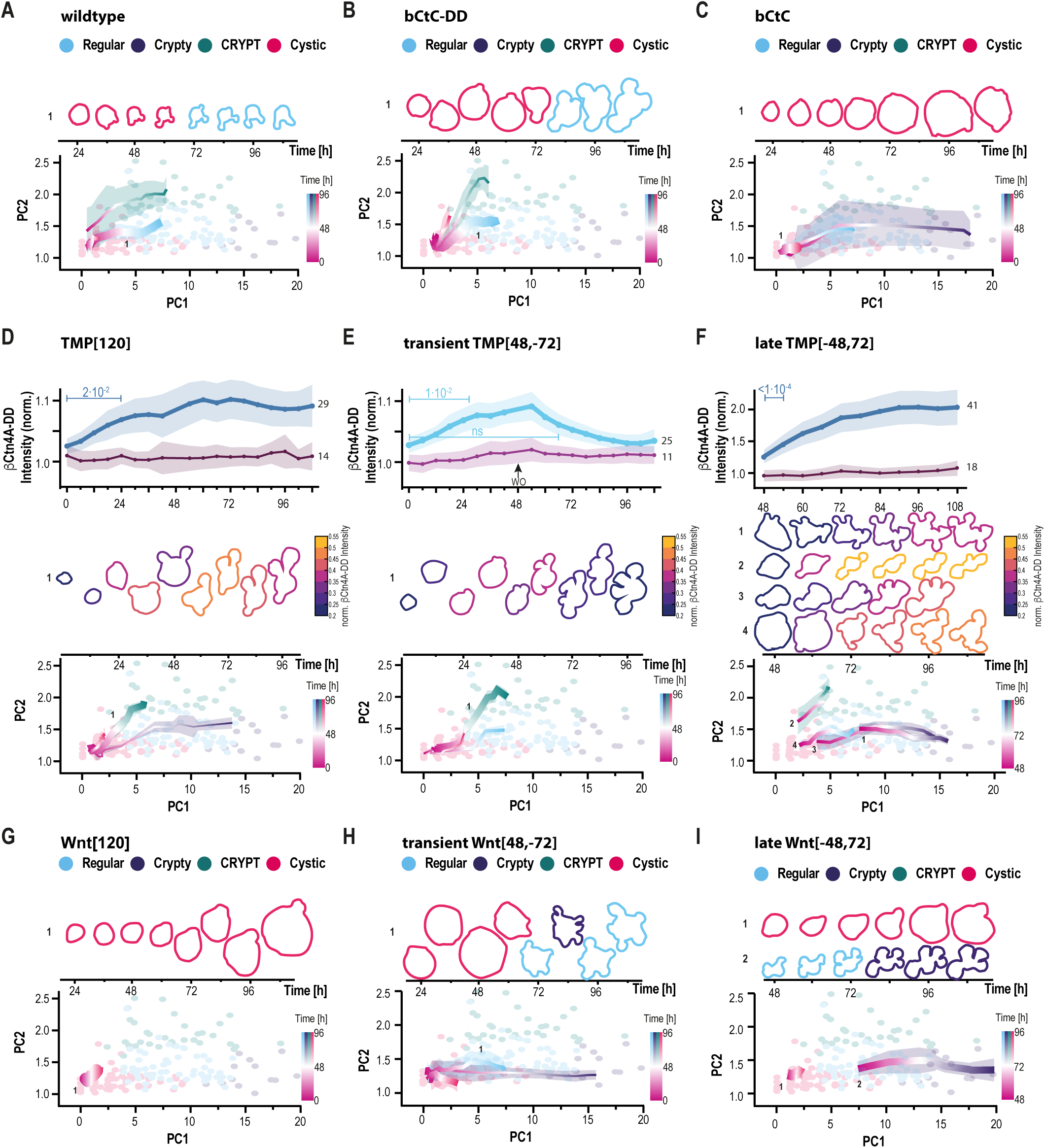
The relation between oncoprotein expression and developmental trajectories. **A.** Top: MIP contours of a single developing wt mSIO most proximal to the main trajectory. Bottom: average morphodynamic trajectories in principal component space (PC1, PC2) grouped by end-state Mclass at 96 h. Line color: Development over time from 0 to 96 h colored by end-stage Mclass: blue: ‘Regular’, violet: ‘Crypty’, green: ‘CRYPT’, red: ‘Cystic’. Line thickness: Mclass fraction at 96 h. N=4; n= 55(10/8/18/19). **B.** MIP contours and average morphodynamic trajectories as in (A) for bCtC-DD Vehicle[120] mSIOs. N=6; n= 71(15/14/19/15/3/5). **C.** MIP contours and average morphodynamic trajectories as in (A) for bCtC mSIOs. N=4; n=40(11/9/11/9). **D.** Top: relative βCtn4A-DD intensity (Intensity / Mean (Vehicle)) for mSIOs with significant (blue) or without (plum) response to TMP[120]. TMP[120]: 120 h exposure to 10 µM TMP. Mean±95%CI; N=2, n=29/14. p-values: two-way ANOVA; Dunnett’s multiple comparisons test. Center: MIP contours of a single developing mSIO most proximal to the main trajectory. Color: Min-Max normalized relative βCtn4A-DD intensity. Bottom: Average morphodynamic trajectories of mSIOs with response to TMP[120] with color indicating development over time from 0 to 96 h and line thickness indicating Mclass fractions as in (A). N=2, n=29. **E.** Top: Relative βCtn4A-DD intensity (Intensity / Mean (Vehicle)) for mSIOs with significant (light blue) or without (purple) response to transient TMP[48,-72]. 1 TMP[48,-72]: 48 h exposure to 10 µM TMP followed by washout and further growth for 72 h in ENR medium. Mean±95%CI; N=2, n=25/11. p-values: two-way ANOVA; Dunnett’s multiple comparisons test. Contours and morphodynamic trajectories as in (D). N=2, n=25. **F.** Top: Relative βCtn4A-DD intensity (Intensity / Mean (Vehicle)) for mSIOs with significant (blue) or without (plum) response to late TMP[-48,72]. TMP[-48,72]: 48 h growth in ENR followed by 72 h exposure to 10 µM TMP. Mean±95%CI; N=3, n=41/18. p-values: two-way ANOVA; Dunnett’s multiple comparisons test. Contours and morphodynamic trajectories as in (D), additionally separated by initial Mclass ‘Regular’ (trajectory 1) or ‘Cystic’ (trajectories 2-4) at 48 h. N=3, n=41. **G.** MIP contours and average morphodynamic trajectories as in (A) for bCtC-DD organoids in Wnt[120]. Wnt[120]: 120 h exposure to 50% Wnt. N=3; n=68(27/22/19). **H.** MIP contours and average morphodynamic trajectories as in (A) for bCtC-DD organoids in transient Wnt[48,-72]. Wnt[48,-72]: 48 h exposure to 50% Wnt followed by washout and 72 h further culture in ENR medium. N=3; n=52(24/18/10). **I.** MIP contours and average morphodynamic trajectories as in (A), additionally separated by initial Mclass ‘Regular’ or ‘Cystic’ at 48 h for bCtC-DD organoids in late Wnt[-48,72]. Wnt[-48,72]: 48 h culture in ENR followed by 72 h exposure to 50% Wnt. N=4; n=90(16/22/21/31).

Mclass end-stage distributions of living organoids at 96 h (Fig. EV4C) exhibited a similar pattern to that of fixed mSIOs (Fig. 3F) including the distinct morphological manifestations associated with acutely induced, chronic or Wnt-mediated β-catenin activity. bCtC-DD and wt mSIO exhibited a similar main developmental trajectory towards the ‘Regular’ Mclass (Fig. 4A, B). However, bCtC-DD organoids exhibited an enhanced growth rate resulting in ∼1.6-fold [CI_0.97_:1.2-2.1] increase in the projected area with respect to wt (Fig. EV4B), as well as an average delay of ∼12 h for the onset of crypt formation (Fig. 4A,B, EV4D). Despite these dissimilarities in growth, the transgenic organoids did not exhibit a significant change in end-stage (96 h) Mclass distribution with respect to wt (Fig. EV4C).

bCtC mSIOs stably expressing βCtn4A-Cit oncoprotein mainly exhibited a growing, morphodynamically static state within the ‘Cystic’ Mclass with a ∼1.5-fold [CI_0.97_:1.0-2.3] enhanced projected area with respect to bCtC-DD mSIOs (Fig. 4C, Fig. EV4B,D). This result contrasting the distinct development towards the ‘CRYPT’ morphology in bCtC-DD mSIOs induced by TMP[120] (Fig. EV4E, top left) indicates that oncoprotein-induced aberrant trajectories depend on the developmental stage at which oncoprotein activity manifests.

In order to investigate this further, we related chemically-controlled temporal profiles of βCtn4A-DD oncoprotein levels to morphodynamic trajectories. To address this, developing bCtC-DD mSIOs were subjected to _(i)_continuous TMP[120] with TMP exposure for 120 h, _(ii)_transient TMP[48,-72] with TMP exposure for the initial 48 h followed by 72 h without TMP, or _(iii)_late TMP[-48,72] with TMP exposure after 48 h of development for 72 h, while βCtn4A-DD expression was monitored by widefield fluorescence microscopy (Fig. EV4F). The obtained average mCitrine fluorescence of βCtn4A-DD was then quantified along evolving bCtC-DD mSIO contours.

During TMP[120], 29 out of 43 bCtC-DD mSIOs exhibited a significant increase in βCtn4A-DD oncoprotein 24 h after TMP exposure (Fig. 4D). The majority of these organoids (16/29) exhibited a steep morphodynamic trajectory towards the aberrant ‘CRYPT’ Mclass (Fig.4D, middle, bottom) with minor fractions exhibiting ‘Cystic’ (7/29), ‘Crypty’ (4/29) and ‘Regular’ (2/29) trajectories (Fig.4D, bottom).

Reversal of oncoprotein stabilization during bCtC-DD development in transient TMP[48,-72] was performed by ecDHFR-assisted TMP washout at 48 h after initial TMP addition (Fig. EV4F,G). This caused βCtn4A-DD to rise to significant levels in 25 out of 36 mSIOs within 30 h, to later decline to levels indistinguishable from initial basal levels at 66 h (Fig. 4E, top, middle). The majority of these organoids maintained the aberrant ‘CRYPT’ trajectory (13/25) with minor fractions exhibiting ‘Regular’ (7/25), ‘Cystic’ (4/25) and ‘Crypty’ (1/25) trajectories (Fig.4E). This corroborates that the early oncoprotein induced developmental trajectory defines continuation of the trajectory towards the aberrant phenotype, even in the absence of the initiating oncoprotein activity, likely due to induced alterations in cellular composition and crypt architecture maintaining the aberrant niche. In continuous and transient TMP administration the short ‘Cystic’ trajectory paralleled the steep ‘CRYPT’ trajectory, indicative of a delayed development of smaller mSIO seeds towards ‘CRYPT’.

To substantiate that the initial morphodynamic trajectory defines the oncoprotein-induced aberrant development, bCtC-DD mSIO precursors were cultured for 48 h before addition of TMP (late TMP[-48,72]). The distribution in developmental states among transgenic organoids 48 h after seeding (Hartl *et al*, 2019) was sufficiently diversified to correlate current to future morphology (Fig. EV4H). During TMP[-48,72], 41 out of 59 bCtC-DD mSIOs exhibited a significant rise in βCtn4A-DD oncoprotein levels 6 h after TMP exposure (Fig. 4F; top, middle). The predominant manifestation at 96 h of ‘Crypty’ instead of ‘CRYPT’ as in early TMP exposure of TMP[120] or TMP[48,-72] (Fig. EV4C) indicates that the morphology prior to TMP exposure further affects the subsequent aberrant morphodynamic trajectories. Indeed, mSIOs that had already reached a differentiated ‘Regular’ Mclass, continued along the aberrant ‘Crypty’ trajectory with further development of initial morphological crypt features (Fig. 4F middle; bottom: trajectory 1). Less differentiated ‘Cystic’ Mclass mSIOs developed along three main trajectories in dependence on fine-grained initial morphological features defining the positioning within the ‘Cystic’ Mclass (Fig. 4F; bottom). Those mSIO’s that showed initial ‘CRYPT’-like morphological features such as few crypt precursors and elevated eccentricity would further develop along that ‘CRYPT’ trajectory (Fig. 4F; middle; bottom: trajectory 2). Those mSIO’s with extended crypt precursors regions (bulge) developed far into the ‘Crypty’ Mclass region, possibly by crypt fission, (Fig. 4F; middle; bottom: trajectory 3), while less developed mSIOs with rudimentary crypt precursor areas followed a ‘Regular’ trajectory (Fig. 4F; middle; bottom: trajectory 4).

The irreversible trajectories upon transient βCtn4A-DD oncoprotein induction were in stark contrast to those that occurred upon transient global Wnt exposure (Fig. 4G,H). Continuous Wnt[120] maintained growth of bCtC-DD mSIO precursors within the ‘Cystic’ Mclass (Fig. 4G) with a ∼1.8-fold [CI_0.97_:1.3-2.7] enhanced growth rate with respect to bCtC-DD mSIOs (Fig. EV4B). This indicates that external Wnt maintains an extensively proliferating undifferentiated fetal-like state (Sato *et al*, 2011b) with massive shedding of cells in the mSIO lumen that eventually results in mSIO death (Fig. EV4E; bottom left). However, transient Wnt[48,-72] caused a major fraction of ‘Cystic’ mSIOs to redevelop crypt structures at ∼72 h leading to ‘Regular’ or more extended ‘Crypty’ morphodynamic trajectories (Fig. 4H) with a ∼1.0-fold [CI_0.97_:0.7-1.9] nominal growth rate. During late Wnt[-48, 72], mSIOs in the ‘Cystic’ Mclass maintained their cystic growth, whereas those in the ‘Regular’ Mclass region further developed along the ‘Crypty’ trajectory. In these mSIOs the crypts itself showed cystic growth, expanding to large structures which indicate disturbed differentiation (Fig. 4I, Fig. EV3E bottom, right).

External Wnt exposure thus reversibly generates a fetal state only in presence of Wnt with delayed differentiation after Wnt removal (Fig. 4 G,H; Fig. EV4E). In contrast, early oncoprotein stabilization irreversibly generates an aberrant developmental trajectory (Fig. 4 D,E; Fig. EV4E) which is maintained even when the oncoprotein is only transiently present. This indicates that the prior phenotype affects future development, onset and duration of oncomimetic β-catenin activity thus determine distinct aberrant developmental routes.

### Local oncoprotein activity globally disturbs organoid development

To next test the opto-chemical stabilization of βCtn4A-DD in organoids, transgenic bCtC-DD mSIOs were cultured for 48 h, incubated with NvocTMP-Cl and entirely irradiated to enable βCtn4A-DD stabilization. Therefore, organoids were intensily mechanically disintegrated to generate small crypt fragments and subsequently cultured for 48 h. By this, the majority of mSIOs showed a small, round morphology (Fig. EV4 H; Cystic) at the timepoint of NvocTMP-Cl (10 µM) administration and irradiation of the entire mSIO with 405 nm light for 4 s (10 J/cm²; methods). In 10 out of 15 mSIOs relative βCtn4A-DD fluorescence significantly increased within 3 h after irradiation (Fig. 5A, B top) while the fluorescence fold-change of the activated cell tracker PAmCherry showed an instant increase followed by subsequent reduction over time during mSIO development (Fig. 5A, B bottom). The corresponding βCtn4A-DD to PAmCherry ratio (Fig. EV5A) can be used to correct for intensity changes due to technical reasons (re-focusing) and dilution of both signals over the growth phase, and shows the contrast between the significant increase of the responding mSIO fraction versus the non-responding mSIOs which show signals comparable to the irradiated Vehicle control, indicating an expression level dependency of βCtn4A-DD stabilization. At this developmental stage, 405 nm irradiation of the complete organoid led to elevated cell death and rupture of mSIOs (Fig. EV5B, top), as visible from the drop in surviving mSIOs, and stagnation of growth (Fig. EV5B, bottom), indicating phototoxicity, with mSIOs treated with NvocTMP-Cl being less sensitive.

**Figure 5.**
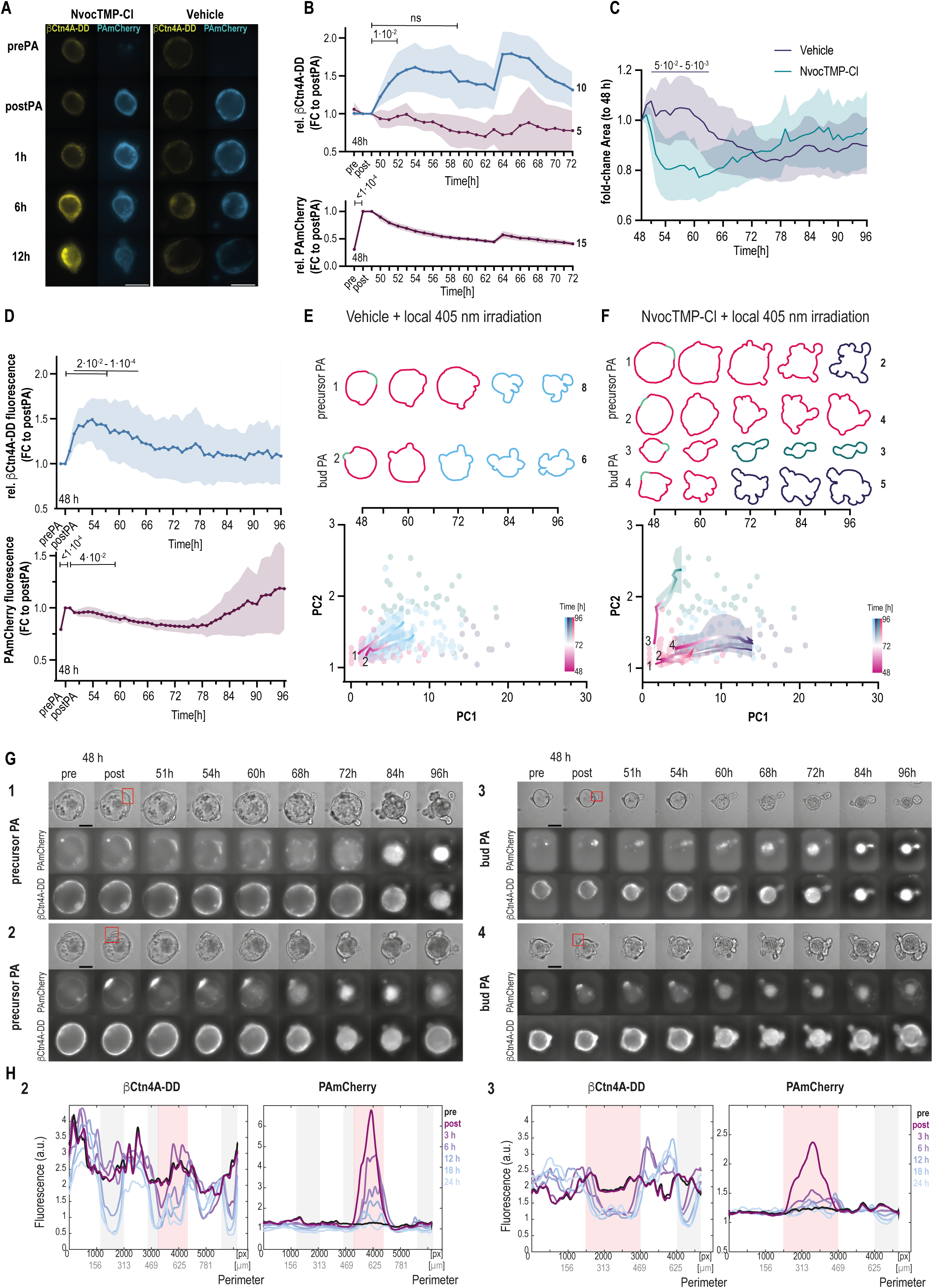
Aberrant development due to local, opto-chemical oncoprotein activity during organoid regeneration. **A.** Representative micrographs of βCtn4A-DD (yellow) and PAmCherry (blue) fluorescence in bCtC-DD mSIOs at indicated timepoints after NvocTMP-Cl (10 µM; left) or Vehicle (0.1% DMSO; right) treatment for 90 min, washout and 405 nm irradiation of the entire mSIO for 4s (10 J/cm^2^). Scale bar: 100 µm. **B.** Top: βCtn4A-DD levels of NvocTMP-Cl (10µM) treated and entirely 405 nm irradiated (10 J/cm^2^) bCtC-DD mSIOs over time relative to Vehicle control (0.1% DMSO; irradiated); normalized to postPA timepoint. mSIOs are separated by high (blue) and low (plum) stabilization of βCtn4A-DD. Mean±95% CI. N=2; n=15. p-values: 2way ANOVA; Dunnett’s multiple comparison test compared to postPA. Bottom: Corresponding PAmCherry fluorescence over time normalized to postPA timepoint. Mean±95% CI. N=2; n=15. p-values: 2way ANOVA; Dunnett’s multiple comparison test compared to postPA. Re-focused at 64 h. **C.** Organoid growth as measured by fold-change of cross-sectional area to 48 h prePA timepoint of mSIO contours from segmented transmission images over time for bCtC-DD Vehicle and NvocTMP-Cl with local irradiation at crypt precursor regions for 4s (10 J/cm^2^). Mean±95% CI. N=5; Vehicle n=16(5/1/2/4/4); NvocTMP-Cl n=16(3/0/2/9/2). p-values: 2way ANOVA; uncorrected Fisher’s LSD. **D.** Top: βCtn4A-DD levels of NvocTMP-Cl (10µM) treated and locally, in crypt precursor regions, 405 nm irradiated (4s; 10 J/cm^2^) bCtC-DD mSIOs over time relative to corresponding locally irradiated Vehicle control; normalized to postPA timepoint. Mean±95% CI. N=5; n=16. p-values: 2way ANOVA; Dunnett’s multiple comparison test compared to postPA. Bottom: Corresponding PAmCherry levels over time normalized to postPA timepoint. Mean±95% CI. N=5; n=16. p-values: 2way ANOVA; Dunnett’s multiple comparison test compared to postPA. **E.** Top: Contours of a single developing Vehicle-treated bCtC-DD mSIO with local 405 nm irradiation (PA) at early precursor (1) or budded (2) crypt regions; most proximal to the main trajectory. Colored by Mclass: blue: ‘Regular’, violet: ‘Crypty’, green: ‘CRYPT’, red: ‘Cystic’. PA region indicated (light green). Bottom: average morphodynamic trajectories in principal component space (PC1, PC2) grouped by PA region (precursor PA: 1; bud PA: 2) and end-state Mclass at 96 h. Line color: Development over time from 48 h to 96 h colored by gradient to end-stage Mclass. Line thickness: Mclass fraction at 96 h. N=5; precursor PA n=8; bud PA n=6. **F.** Top: Contours of a single developing NvocTMP-Cl-treated bCtC-DD mSIO with local 405 nm irradiation (PA) at early precursor (1/2) or budded (3/4) crypt regions; most proximal to the main trajectory. Colored by Mclass: blue: ‘Regular’, violet: ‘Crypty’, green: ‘CRYPT’, red: ‘Cystic’. PA region indicated (light green). Bottom: average morphodynamic trajectories in principal component space (PC1, PC2) grouped by PA region and end-state Mclass at 96 h colored by gradient to end-stage Mclass. Line thickness: Mclass fraction at 96 h. N=5; precursor PA n=6; bud PA n=8. **G.** Micrographs (maximum intensity projections) of developing bCtC-DD NvocTMP-Cl mSIOs with local 405 nm irradiation in Transmission, PAmCherry and βCtn4A-DD pre and post local irradiation (PA; red rectangle) at indicated timepoints for precursor PA (left; 1,2) and bud PA (right; 3,4) corresponding to the contours and trajectories in (F). **H.** Linearized βCtn4A-DD and PAmCherry fluorescence along the developing mSIO contours for NvocTMP-Cl treated examples 2 (precursor PA; left) and 3 (bud PA; right) at indicated timepoints (prePA, postPA, 3 h, 6 h, 12 h, 18 h, 24 h) after PA with normalized region sizes and and matched regions (villus and crypt regions at 18 h after PA). Background color indicates regions: red – PA region; grey – additional crypts.

In response to tissue injury, freshly seeded crypts undergo a transient fetal-like reprogramming characterized by a YAP/TAZ-dependent transcriptional program (Nusse *et al*, 2018; Viragova *et al*, 2024; Yui *et al*, 2018). In this regenerative, fetal-like state, mSIOs have a layer of short cuboidal cells (as in Appendix Fig. S5 A; 60h) which can be distinguished from the tall columnar shape of the villus region in differentiated mSIOs (as in Appendix Fig. S5A; 78h). These two distinct states during organoid development enable analysis of locally induced oncoprotein activity during homeorhesis (regenerative, fetal-like mSIO) versus homeostasis (differentiated mSIO).

In order to study how the locally induced oncoprotein activity in mSIOs may alter the development of the entire organoid, we first targeted undifferentiated mSIOs in the fetal-like state during homeorhesis.

In this state, distinct developmental states of the crypt precursor regions can be distinguished. Bulge regions, as earlier crypt-precursor states, and buds, as later crypt-precursor states, develop during early crypt formation (Appendix Fig. S5A) (Hartl *et al*., 2019; Xue *et al*, 2025; Yang *et al*, 2021). The bulge precursor regions (precursor PA), typically already contain few paneth cells and have columnar shape compared to the thin cell layer of the surrounding fetal-like tissue. The budded states (bud PA) typically show initiated apical constriction of the developing crypt (Xue *et al*., 2025; Yang *et al*., 2021) and a prototypical contrast of βCtn4A-DD fluorescence between the developing crypt and villus regions. This contrast is also observed in end-stage mSIOs after 96 h of growth, showing a ratio of 1.7±0.4 for the β-catenin signal in the Aldolase B-positive villus region compared to the crypts (Fig. EV5C), likely originating from the distinct levels of E-Cadherin and thus distinct levels of β-catenin in adherens junctions.

Initial global analysis showed an altered growth behavior upon local βCtn4A-DD stabilization in crypt precursor regions of developing mSIOs exhibiting a significantly reduced fold-change of the cross-sectional Area (FC Area) around 3-15 h after irradiation compared to the locally irradiated vehicle control (Fig. 5C).

The vehicle control showed a typical lumen deflation (Xue *et al*., 2025) after 60 h of culture, typically representing the switch from regenerative/fetal-like to differentiated state, followed by a phase of organoid homeostasis and crypt growth. In contrast, NvocTMP-Cl treated and locally irradiated mSIOs showed an early abrupt deflation followed by gradual growth. This phase was characterized by inflation – deflation cycles of the lumen, as indicated by the spikes in the growth curve, as well as crypt budding, fission and growth. Relative βCtn4A-DD fluorescence measured over the whole mSIO contour increased significantly within 4 h after local irradiation and decreased back to non-significant levels after 16 h (Fig. 5D, top). Likewise, PAmCherry fluorescence – co-activated in the instance of NvocTMP-Cl uncaging – decreased due to dilution during proliferation and cell shedding resulting in significantly reduced levels 11 h after irradiation (Fig. 5D, bottom). Compared to the locally irradiated vehicle, the fold-change of the βCtn4A-DD fluorescence was significantly increased 1-9 h after irradiation (Fig. EV5 D, left).

Notably, morphodynamic trajectories of the locally irradiated vehicle control showed a development towards the ‘Regular’ Mclass irrespective of the developmental state of the irradiated crypt precursor region (Fig. 5E). Thus, in developing mSIOs with clear crypt precursor regions the local irradiation with 405 nm for 4 s (10 J/cm²) does not lead to aberrant development or increased organoid damage. Thereby, this developmental state can be used to study consequences of oncomimetic β-catenin activity during homeorhesis. Local opto-chemical βCtn4A-DD stabilization in both developmental states of crypt precursor regions, precursorPA (Fig. 5F 1,2) and budPA (Fig. 5F 3,4), led to aberrant development of the whole organoid characterized eithter by enrichment or enlargement of crypts or crypt-like regions and resulting ‘Crypty’, ‘Cystic’ or ‘CRYPT’ Mclasses.

As observed for global βCtn4A-DD stabilization by TMP, trajectories of mSIOs following local βCtn4A-DD stabilization were not obviously generalizable and depended on multiple factors such as the developmental state of the activated crypt precursor region, number and developmental state of additional crypt precursor regions, and the overall developmental state of the organoid. Sustained βCtn4A-DD stabilization in budPA mSIOs following the ‘CRYPT’ trajectory most likely reflects reduced proliferation compared to mSIOs developing along the ‘Crypty’ trajectory rather than being the primary cause of trajectory divergence (Fig. EV5 D, right).

We continued to investigate the local development of βCtn4A-DD and corresponding PAmCherry fluorescence in representative mSIOs of aberrant developmental trajectories (Fig. 5F-H, EV5E) and vehicle controls (Fig. 5E, EV5F) over 24 h.

The instantaneous increase of PAmCherry fluorescence (Fig, 5H, EV5E, F– light red region) in the irradiated regions (Fig. 5G, EV5 F top – red rectangle;) was detected for precursorPA and budPA in NvocTMP-Cl treated mSIOs and the vehicle control. Due to the typically lower βCtn levels in crypts compared to the villus, βCtn4A-DD stabilization can be primarily perceived as peaks moving over time in the villus regions flanking the irradiated region (Fig. 5H, EV5E, Appendix Fig. S5 B,C). The fluorescence intensity villus to crypt relation reached a ratio > 2 within 6 h after irradiation (Fig. EV5G). In comparison, βCtn4A-DD levels in the vehicle control showed prototypical villus to crypt ratios of ∼1.6 around 18-24 h after PA (Fig. EV5G) during differentiation and establishment of the ‘Regular’ Mclass morphology (Fig. EV5F, G, Appendix Fig. S5 D). In addition, developing crypts show a significantly increased activated/non-activated βCtn4A-DD fluorescence upon precursorPA (Fig. EV5 H). Irradiated crypt precursor regions in the NvocTMP-Cl treated mSIOs also showed a significant increase of cumulative βCtn4A-DD levels in the irradiated regions compared to the vehicle control. Furthermore, upon budPA NvocTMP-Cl-treated mSIOs showed a slightly elevated crypt growth than the vehicle control (Fig. EV5 I). These results highlight that locally induced oncogenic β-catenin activity in crypt precursor regions can induce global aberrant development in regenerating intestinal tissues which is characterized by expansion and enrichment of the crypts.

### Local oncoprotein activity in mature crypts disrupts organoid homeostasis

To further study if oncogenic β-catenin activity impacts the tissue morphogenesis not only during homeorhesis but also during the more stable state of tissue homeostasis, mature mSIOs with a well-defined crypt-villus structure were targeted at early (small) or late (big) crypts.

In this state of differentiation, local opto-chemical βCtn4A-DD stabilization in crypts led to increased growth as shown by the fold-change of the overall organoid area compared to the vehicle control (Fig. 6A). Furthermore, the global βCtn4A-DD fluorescence of NvocTMP-Cl relative to the vehicle control was significantly increased during 2-7 h after irradiation, normalized signals for both conditions showed an elevated plateau in the NvocTMP-Cl condition compared to the stable mean of the vehicle control (Fig. 6B; Fig EV6B).

**Figure 6.**
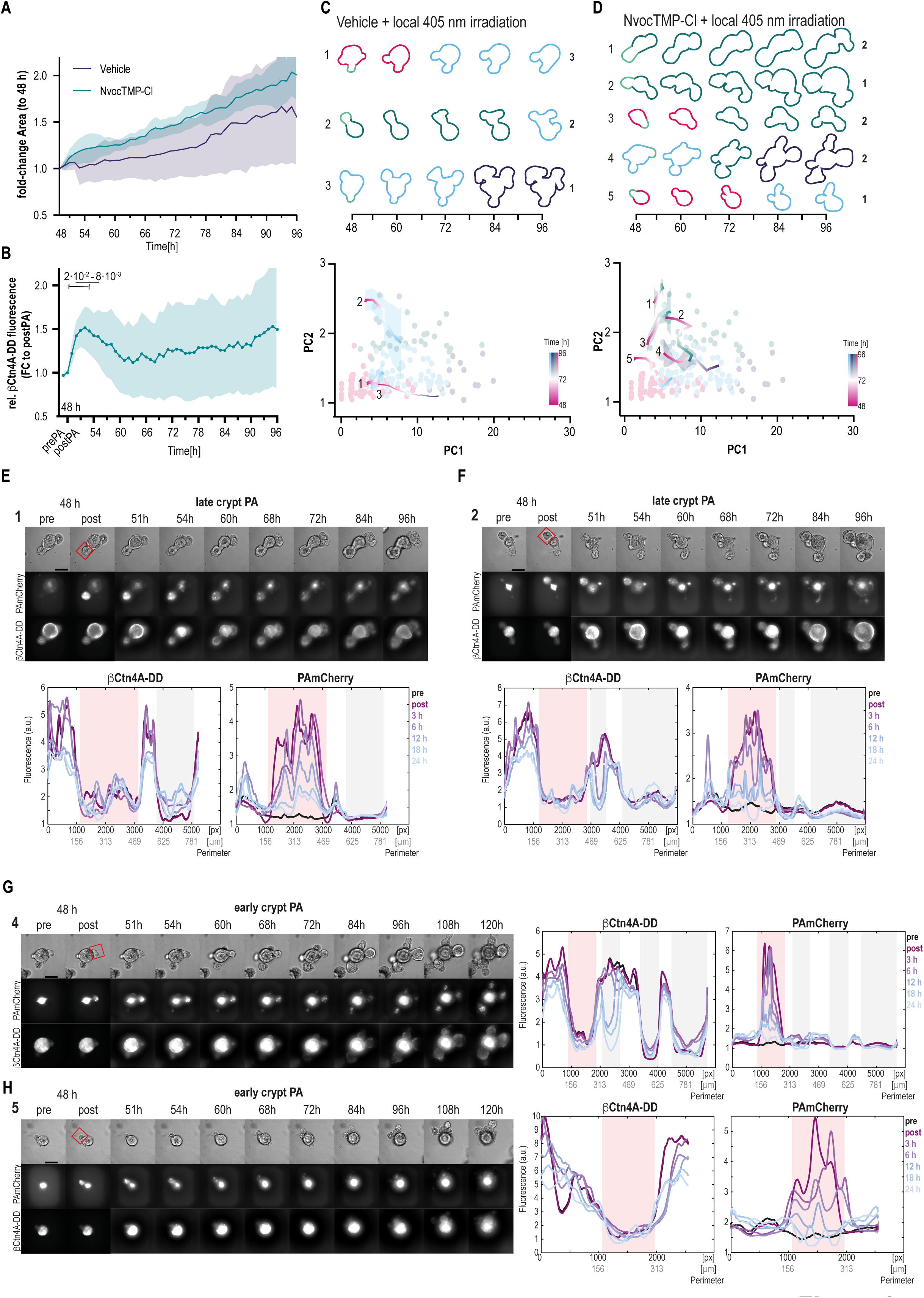
Aberrant organoid development after local, opto-chemical oncoprotein stabilization during homeostasis. **A.** Organoid growth as measured by fold-change to 48 h prePA timepoint of cross-sectional area of mSIO contours from segmented transmission images over time for bCtC-DD Vehicle (0.1% DMSO) and NvocTMP-Cl with local irradiation at early and late crypts of differentiated mSIOs for 4s (10 J/cm^2^). Mean±95% CI. N=5; Vehicle n=6; NvocTMP-Cl n=8. **B.** Relative expression of βCtn4A-DD NvocTMP-Cl (10µM) to Vehicle (0.1% DMSO) treated and locally, in crypt regions, 405 nm irradiated (10 J/cm^2^) differentiated bCtC-DD mSIOs over time; normalized to postPA timepoint. Mean±95% CI. N=5; n=8. p-values: 2way ANOVA; Dunnett’s multiple comparison test compared to postPA. **C.** Top: Contours of a single differentiated bCtC-DD Vehicle mSIOs with local 405 nm irradiation (PA) at crypt regions; most proximal to the main trajectory. Colored by Mclass: blue: ‘Regular’, violet: ‘Crypty’, green: ‘CRYPT’, red: ‘Cystic’. PA region indicated (light green). Bottom: average morphodynamic trajectories in principal component space (PC1, PC2) grouped by crypt (early, late) and end-state Mclass at 96 h. Line color: Development over time from 48 h to 96 h colored by gradient to end-stage Mclass. Line thickness: Mclass fraction at 96 h. N=5; n=6. **D.** Top: Contours of a single differentiated bCtC-DD NvocTMP-Cl (10µM) mSIOs with local 405 nm irradiation (PA) at crypt regions; most proximal to the main trajectory. Colored by Mclass: blue: ‘Regular’, violet: ‘Crypty’, green: ‘CRYPT’, red: ‘Cystic’. PA region indicated (light green). Bottom: average morphodynamic trajectories in principal component space (PC1, PC2) grouped by crypt (early, late) and end-state Mclass at 96 h. Line color: Development over time from 48 h to 96 h colored by gradient to end-stage Mclass. Line thickness: Mclass fraction at 96 h. N=5; n=8. **E.** Top: Micrographs (maximum intensity projections) of exemplary developing bCtC-DD NvocTMP-Cl mSIOs with local 405 nm irradiation in Transmission, PAmCherry and βCtn4A-DD pre and post local irradiation (PA; red rectangle) at indicated timepoints for late cryptPA (trajectory 1). Bottom: Linearized βCtn4A-DD and PAmCherry fluorescence along the developing mSIO contours for the NvocTMP-Cl treated examples 1 (late crypt PA) at indicated timepoints (prePA, postPA, 3 h, 6 h, 12 h, 18 h, 24 h) after PA with normalized and matched region sizes (villus and crypt regions at 18 h after PA). Background color indicates regions: red – PA region; grey – further crypts. **F.** Same as in E for the exemplary developing bCtC-DD NvocTMP-Cl mSIOs with late cryptPA (trajectory 2). **G.** Left: Micrographs (maximum intensity projections) of exemplary developing bCtC-DD NvocTMP-Cl mSIOs with local 405 nm irradiation in Transmission, PAmCherry and βCtn4A-DD pre and post local irradiation (PA; red rectangle) at indicated timepoints for early cryptPA (trajectory 3). Bottom: Linearized βCtn4A-DD and PAmCherry fluorescence along the developing mSIO contours for the NvocTMP-Cl treated examples 4 (early crypt PA) at indicated timepoints (prePA, postPA, 3 h, 6 h, 12 h, 18 h, 24 h) after PA with normalized and matched region sizes (villus and crypt regions at 18 h after PA). Background color indicates regions: red – PA region; grey – further crypts. **H.** Same as in G for the exemplary developing bCtC-DD NvocTMP-Cl mSIOs with early cryptPA (trajectory 5).

The morphodynamic trajectories showed that irradiated vehicle-treated mSIOs mainly developed towards the ‘Regular’ morphology (Fig. 6C) after irradiation of early, smaller (trajectory 1) and late, bigger (trajectory 2) crypts. In contrast to the elevated crypt fission during homeorhesis, NvocTMP-Cl-mediated βCtn4A-DD stabilization in crypts of mature mSIOs mainly led to aberrant development of the present crypts, also resulting in crypt fission in few examples (Fig. 6D). This development was comparatively faster in mSIOs with activated later crypts (Fig. 6D, trajectories 1&2; Fig. 6E,F) compared to the slower manifestation in mSIOs with activated early crypts (Fig. 6D, trajectories 3-5; Fig. 6G,H, Fig. EV6 D). Interestingly, in 5 out of 8 mSIOs not only the activated crypts showed an abnormal development following the βCtn4A-DD stabilization but also neighboring, non-activated crypts developed aberrantly and initiated a separation from the original mSIO (Fig. 6 E,G; trajectories 1,4&5) and independently grew out new crypt-villus structure of an individual mSIO. In the vehicle control only 1 out of 6 mSIOs showed an initiated separation of the activated crypt, which fused again with the villus body (Fig. EV6 F, Video 2)

The linearized fluorescence signals of βCtn4A-DD showed significantly increased level of cumulative βCtn4A-DD fluorescence compared to the vehicle and slightly higher levels in the activated crypt compared to non-activated crypts of the same mSIO (Fig. 6 E-H, Fig EV6 E-G, Fig. EV6 I). Furthermore, in NvocTMP-Cl-treated mSIOs the activated crypts showed a tendency of elevated growth compared to the vehicle control (Fig. EV6 H). However, neither βCtn4A-DD nor PAmCherry fluorescence along the mSIO contours showed a clear spreading of the fluorescence signals into villus regions flanking the targeted crypts (Fig. 6 E-H) for mSIOs exposed to NvocTMP-Cl while the vehicle control (Fig. EV6 E,F) showed a moving peak for PAmCherry in the adjacent villus regions.

These results showed that the oncogenic activity of βCtn4A-DD in crypts disturbs also tissue organization during homeostasis leading to aberrant development of the activated crypts and destabilization of the non-activated crypts inducing a separation from the tissue. This might indicate that the cell migration from crypts to the villus region is disturbed upon βCtn4A-DD stabilization.

## Discussion

In this study, we developed a chemical-(opto)genetic system to induce oncomimetic β-catenin activity globally by TMP or locally by NvocTMP-Cl, respectively. Using this approach, we investigated how oncogenic β-catenin activity can affect intestinal tissue transformation and addressed the impact of developmental state, timing and reversibility as well as region of activation and the propagation of the oncogenic β-catenin activity.

In particular, one of the unique features of this chemical-(opto)genetic system is to enable real-time visualization of early tissue transformation events, capturing the interaction of adjacent wildtype cells upon local oncogenic signaling, which was previously not possible. Earlier applications of the photoactivatable NvocTMP-Cl dimerizer system were limited to fast-acting processes in the seconds/minutes scale (Chen *et al*, 2018; Chen & Wu, 2018). The re-engineering in this study through integration of the compound-retaining Halo-3xNLS and the combination with the DD expanded its capability to achieve rapid, sustained stabilization of the target protein over days, a timeframe which allows the study of long-term processes such as development and tumorigenesis. Furthermore, this approach can be adapted to elucidate other oncogenic mutations, especially in combination with a nuclear function, and provides a versatile platform for spatiotemporal-control of transcription factors or epigenetic regulators, a task which until recently remains technically challenging (Zhu *et al*, 2023).

We applied this approach in the well-established model system of small intestinal organoids (Clevers, 2026) to mimic initial changes caused by APC loss-of-function mutations in the early-phase of CRC development. In this model, eithter freshly seeded intestinal stem cells or crypts initially recapitulate a phase of tissue repair and present fetal-like characteristics (Fey *et al*, 2024; Nusse *et al*., 2018; Viragova *et al*., 2024; Yui *et al*., 2018). This phase of organoid re-development after injury is optimal to test robustness of the tissue homeorhesis towards oncoprotein activity (Ray, 2021; Waddington, 2014). In later stages the YAP signaling dominated fetal-like organoid state changes into a differentiated/mature Wnt signaling dominated state of tissue homeostasis, resembling an adult tissue (Fey *et al*., 2024; Serra *et al*., 2019). In this phase, local induction of oncogenic β-catenin activity in crypts is expected to mimic APC loss-of-function mutations and allows to study the interaction of a potentially transformed crypt with the normal surrounding tissue during early tissue transformation.

Investigating the capability of oncomimetic β-catenin mutations, the continuous expression of βCtn4A-Cit in mSIOs led to a cystic development. This phenotype is similar to the spheroid formation described for small intestinal organoids harboring inactivating APC mutations (Farrall *et al*, 2012; Huels *et al*., 2015; Sato *et al*., 2011a), inactivating GSK3 mutations (Andersson-Rolf *et al*, 2017) or homozygous activating β-catenin mutations (Farrall *et al*., 2012; Huels *et al*., 2015), closely resembling the previously described spheroid-like growth upon external Wnt-signaling activation. However, after 2-3 days of growth the organoids differentiated as recognized by the switch of cell shape from the smooth, thin-walled fetal-like spheroid to the thicker columnar cell layer and a ruffled appearance (Video 1). In contrast to APC^Min/Min^ spheroids with a sparse, homogenous Paneth cell distribution (Langlands *et al*, 2018) the continuous expression of βCtn4A-Cit resulted in a shift towards the Paneth cell lineage. These changes resemble the elevated proliferation, crypt hyperplasia and commitment towards the Paneth cell lineage in APC-deficent mouse models (Andreu *et al*., 2005; Andreu *et al*, 2008; van Es *et al*, 2005). This phenotype is commonly described as crypt precursor phenotype and indicates a strong shift towards less-differentiated, progenitor-like cell states (Jarde *et al*, 2013).

The global, acute stabilization of βCtn4A-DD during organoid re-development from seeded crypts resulted in crypt enlargement and expansion of the stem cell zone, which resembles a state similar to the early manifestation of the crypt progenitor phenotype in βCatenin^+/Δex3^ intestine (Huels *et al*., 2015) and of doxycycline induced TetOΔN89 - βCatenin4A expression in organoids (Jarde *et al*., 2013), but differs from the spheroids published in another doxycycline inducible model TetON-βCatenin4A model which reported the spheroid-like growth similar to APC KO organoids (Farrall *et al*., 2012; Riemer *et al*, 2017). In contrast to these publications, the mSIOs in our study were always cultured in presence of R-spondin which promotes and amplifys endogenous Wnt-signaling from Paneth cell-secreted Wnt (de Lau *et al*, 2011). Furthermore, consequences of oncogenic β-catenin expression have been shown to be dose dependent and E-cadherin as binding partner in adherens junctions can act as a buffer limiting transformation potential (Huels *et al*., 2015).

In our study, the TMP-induced total relative β-catenin protein levels were increased more than two times compared to the vehicle control and even compared to the stable expression of bCtn4A in bCtC organoids. Thus, expression levels only cannot explain the distinct phenotypic outcomes between our study and others, which showed mRNA expression levels with fold-changes of more than thousands (Huels *et al*., 2015; Jarde *et al*., 2013). However, these findings suggest that the outcome of oncogenic β-catenin activity is highly context- and dose dependent.

In previous studies, the mouse intestinal tissue was able to re-establish a normal tissue organization after APC restoration (Dow *et al*, 2015) or terminated expression of mutated β-catenin (Jarde *et al*., 2013) over weeks. In this study we targeted the reversibility of the organoids developmental trajectories during re-development after injury. While an early removal after 24 h appears to result in development of normal organoid phenotypes (Farrall *et al*., 2012), the removal at later timepoints like 48 h results in a conserved aberrant trajectory towards organoids with enlarged crypts similar to the tissue crypt precursor phenotype.

This difference in reversibility might be due to the timepoint during the organoid re-development. In the average organoid population redeveloping after seeding single crypts, the 48 h timepoint is much closer to the switch between the fetal-like state and the Wnt-dominated differentiated state than 24 h which is during the YAP-dominated fetal-like state (Serra *et al*., 2019). Regarding the timing, organoids showed reversibility upon transient external Wnt3a stimulation for 48 h with re-estabilishment of budding structures after Wnt removal, a well-known behaviour utilized during organoid transfection and generation of transgenic organoid lines (Heinz *et al*., 2020). In contrast to wildtype β-catenin which is rapidly degraded within hours after Wnt removal (Munemitsu *et al*, 1995), the TMP-stabilized βCtn4A-DD levels decrease back to basal levels within 18 h after ecDHFR-assisted TMP washout. Another possible explanation might be the abundance of βCtn4A-DD in adherens junctions, which can on the one hand act as a reservoir for TMP-stabilized βCtn4A-DD (Kam & Quaranta, 2009) and on the other hand alter mechanical characteristics of cells and their behaviour in migration and wound healing (Leng *et al*, 2020; Tsai & Galko, 2019).

Importantly, the response to oncogenic signaling depended strongly on the developmental state of the organoid. Consistent with this, the late global exposure to Wnt or stabilization of oncogenic β-catenin showed a clear dependency on the organoids developmental state at onset. In case of Wnt exposure, fetal-like spheroids and organoids with budded crypt precursor regions showed two distinct trajectories of cystic growth and aberrant crypt development, respectively.

This initial state dependency was even more pronounced upon late global induction of oncogenic β-catenin activity by addition of TMP after 48 h of growth. This results in aberrant development characterized by crypt enlargement and/or crypt fission which might be induced by oncogenic β-catenin activity-mediated restructured crypt environment (Song *et al*, 2014) leading to enrichment and altered positioning of Paneth cell clusters which are reported to be a driver of crypt fission when flanking LGR5^+^ intestinal stem cell clusters (Langlands *et al*, 2016).

Cellular migration and positioning along the crypt are controlled by differential adhesion and activation of the EphB receptors and ephrinB ligands, which is mediated by β-catenin and TCF (Batlle *et al*, 2002). Simulations have shown that a loss of this differential adhesion leads to random distribution of Paneth cells (Wong *et al*, 2010). This model fits to our results of global TMP-mediated oncogenic β-catenin activity, which showed altered positioning of Paneth cells and early characteristics of the crypt precursor phenotype, but lacked a significant transformation towards polyp or early adenoma features. A reason for that might be the global oncogenic β-catenin activity, which diminishes differences in tissue regions and thus protects the tissue from global transformation.

In contrast, upon local oncogenic β-catenin activation the activated tissue can establish distinct features than the surrounding wildtype tissue. Intrestingly, activation of oncogenic β-catenin in the developing crypt precursor regions of undifferentiated fetal-like organoids also led to crypt enrichment by fission. During this development the organoids performed multiple lumen inflation–deflation cycles. This alternative way of crypt fission is described as a mechanosensitive Piezo-channel dependent mechanism by which the stretch – relaxation cycles lead to separation of stem cell clusters and subsequent budding upon deflation (Tallapragada *et al*, 2021). During homeorhesis, the local oncoprotein activity thus globally resulted in an aberrant phenotypic change.

*In vivo*, these conditions of tissue repair might arise after damages by irradiation, chemotherapy, inflammation or penetrating trauma where the resulting regeneration is characterized by transient activation of fetal programs (Fey *et al*., 2024).

However, CRC is postulated to arise in situations of tissue homeostasis during adulthood from single heterogenous stem cell acquiring a second mutation and propagates bottom-up from the affected crypt base taking over the entire crypt (Huels & Sansom, 2015; Vermeulen *et al*, 2013). In mature organoids, representing the state of tissue homeostasis, the perturbation by local oncogenic β-catenin activation in crypts results in aberrant crypt growth characterized mainly by enlargement of the activated crypt in combination with the trend of initiated crypt separation from the villus for the non-activated crypts.

These observations revealed that the activated, altered crypts disturbed the overall tissue organization and prevented cells from differentiation/migration into the villus, which partially recapitulates the immediate effects of APC loss-of-function reported by previous mouse model studies and correlates with the *in vivo* observations of depletion of mature enterocytes over the malignancy continuum of CRC development (Andreu *et al*., 2005; Becker *et al*, 2022; Sansom *et al*, 2004). To our knowledge, this is the first time this aberrant behavior has been recapitulated in organoids, highlighting the efficacy and innovation of our oncomeme system to recapitulate the early events during oncogenic tissue transformation.

Inspired by the Mem theory from Richard Dawkins (Dawkins, 1976) which describes the phenomenon of evolution-like propagation of cultural ideas from mind to mind or person to person by imitation/mimesis, we coined the term Oncomeme. It describes the ability of a local oncogenic altered unit to influence and alter the surrounding similar units not expressing the causal oncoprotein.

A similar behaviour was described for paracrine signaling between tumoroids and wildtype organoids (Jacquemin *et al*, 2022) which results in tumoroid-like development of the surrounding healthy organoids. It is now further supported by our findings of the spreading of oncogenic behavior from transformed crypt to other crypts of the same tissue.

## Materials and Methods

### Mouse strains

Animal experiments and husbandry were performed according to the German Animal Welfare guidelines and approved by the Landesamt für Natur, Umwelt und Verbraucherschutz Nordrhein-Westfalen (State Agency for Nature, Environment and Consumer Protection of North Rhine-Westphalia). The female C57/BL6 mouse used to generate organoids for this study was at age 23 weeks. The animals were maintained under a 14-hour light/10-hour dark cycle with free access to food and water. Male mice were kept individually, whereas the female mice were housed in groups of up to four per cage.

### Plasmids and Cloning

In order to generate vector plasmids for stable cell line and organoid line generation, plasmid fragments were combined by traditional cloning (NEB restriction enzymes and Quick Ligase) and ligation independent cloning (NEB T4 DNA Polymerase) and cloned into a vector backbone containing PiggyBac transposition sites (Wang *et al*, 2008). Stably transfected lines were created by co-transfection with the CAG-pBASE (Wang *et al*., 2008) transposase and subsequent selection with puromycin.

The final pPB-bCtCDD-PCh-Hal plasmid (12.466 kb) is a combination of three fragments, separated by P2A-T2A ribosomal skipping sites (Donnelly *et al*., 2001a; Donnelly *et al*., 2001b; Liu *et al*., 2017; Pan *et al*., 2017):

- bCtCDD: fused sequences of β-catenin S33A/S37A/T41A/S45A (Liu *et al*., 2002) (bCt), mCitrine (C) and ecDHFR N18T/A19V destabilization domain (DD) (Iwamoto *et al*., 2010). β-catenin S33A/S37A/T41A/S45A was generated by mutation of the mouse β-catenin wildtype sequence from a β-catenin-mCitrine plasmid (Prost, 2015) using NEB Q5® Site-Directed Mutagenesis Kit. ecDHFR N18T/A19V was extracted from pBMN-YFP-DHFR(DD) which was a gift from Thomas Wandless (Iwamoto *et al*., 2010) (addgene # 29326).
- PCh: PAmCherry from pPAmCherry1-N1 was a gift from Vladislav Verkhusha (Subach *et al*., 2009)(addgene # 31928).
- Hal: fused sequences of HA-tag, Halo-tag and 3x NLS (SV40 nuclear localisation sequence). The pPB-bCtCDD-Hal plasmid, lacking PAmCherry was an intermediate result of pPB-bCtCDD-PCh-Hal cloning. The pPB-bCtC is the combination of β-catenin S33A/S37A/T41A/S45A and mCitrine as fusion protein. The pPB-TOPmCherry transcriptional reporter is a modification of M50 Super 8x TOPFlash which was a gift from Randall Moon(Veeman *et al*., 2003) (addgene # 12456).

### Reagents

#### TMP

Trimetroprim (T7883; Sigma-Aldrich);

#### NvocTMP-Cl

custom synthesis by ChiroBlock GmbH (Bitterfeld-Wolfen, Germany) chemical name: (4,5-dimethoxy-2-nitro-phenyl)methyl N-[4-amino-5-[[4-[2-[2-[2-[2-[2-(6-chlorohexoxy)-ethoxy]ethoxy]ethoxy]ethoxy]ethoxy]-3,5-dimethoxy-phenyl]methyl]pyrimidin-2-yl]carbamate; C39H56ClN5O14; M=854.36 g/mol

### Preparation of recombinant ecDHFR

The ecDHFR (*E. coli* Dihydrofolate reductase) coding plasmid pET15b His6-ecDHFR (WT) was a gift from Thomas Wandless (Addgene plasmid # 73188). The recombinant ecDHFR was expressed in E. coli BL21 DE3 RIL (CN: 230245, Agilent, Santa Clara, CA, United States) after induction with 0.2 mM IPTG over night at 18 °C in LB medium. Cell pellets were solved in buffer containing 50 mM NaHPO4, 300 mM NaCl, 0.1 mM β-mercaptoethanol and 10 mM imidazole, pH 8 and lysed by two passes through an Emulsiflex C5 (Avestin, Mannheim, Germany). Lysates was clarified by centrifugation at 48,000 × g in an A27-8 × 50 rotor (Sorvall/Thermo Fisher Scientific) for 45 min. The recombinant protein was purified from the supernatant via Ni-NTA Superflow (Qiagen). The column was washed with 50 mM NaHPO4, 300 mM NaCl, 35 mM imidazole and eluted with additional 500 mM imidazole. The eluate was supplemented with 2-4 mg of Thrombin protease (Cytiva; GE27-0846-01) and dialyzed over night against 2 L of 50 mM Tris pH 8, 100 mM NaCl, 1 mM EDTA, 0.1 mM β-mercaptoethanol. The cleaved protein was adjusted to 10 mM imidazole followed by another pass through the Ni-NTA column. Subsequently size exclusion chromatography was performed on a HiLoad 26/600 Superdex 75 pg column (GE Healthcare) in 50 mM Hepes pH 7.5, 200 mM NaCl, 2 mM DTT. Protein concentration was determined by absorption at 280 nm and aliquots were frozen in liquid nitrogen and stored at -70 °C.

### Cell culture

HeLa cells (ACC-57; Leibniz Institute DSMZ-German Collection of Microorganisms and Cell Cultures GmbH) were cultured in DMEM (PAN Biotech), supplemented with 10% fetal bovine serum (FBS) (PAN Biotech), 2 mM glutamine (PAN Biotech) and 1% Non-Essential Amino Acids (PAN Biotech) (DMEM growth media) at 37 °C with 5% CO_2_ and regularly tested for mycoplasma contaminations using MycoAlert Mycoplasma detection kit (Lonza). Stable transgenic HeLa cell lines were generated via transfection with Fugene 6 (Promega) with a 1:3 DNA to Fugene6 volume ratio. Transgenic cells were isolated based on their puromycin resistance (2 µg/mL) and their mCitrine fluorescence by fluorescence activated cell sorting (FACS) (Aria Fusion Flow Cytometry System, BD Bioscience).

### Live cell imaging of HeLa cells

To investigate the global, chemical stabilization of βCtn4A-DD, HeLa cells stably transfected with pPB-bCtC-DD were seeded into 8-well Lab-Tek chambers (Thermo Fisher Scientific). After 24 h cells were exposed to DMSO as Vehicle or TMP and imaged for 48 h with an Olympus IX81 inverted microscope (Olympus Life Science) containing a MT20 illumination system and incubation chamber at constant 37 °C and 5% CO_2_. Micrographs were acquired every 30 min using a 4x/0.16 NA or 20x/0.75 NA air objective and an ORCA-ER CCD digital camera (Hamamatsu Photonics, Japan). The washout of TMP was performed between the imaging intervals using DMEM growth media. The mean fluorescence of every 32-bit image was obtained after background correction and plotted as mean ± standard error of the mean (SEM) of four different fields of view per condition.

### Opto-chemical stabilization of βCtn4A-DD in HeLa cells

For testing the opto-chemical stabilization of βCtn4A-DD, HeLa cells were cultured for 24 h before incubation with 10 µM NvocTMP-Cl (ChiroBlock GmbH) for 1 h at 37°C. Excess NvocTMP-Cl was removed by washing three times with DMEM growth media. Uncaging of NvocTMP-Cl was performed using a Leica TSC SP8 microscope (Leica Microsystems, Wetzlar, Germany) with incubation chamber and irradiation with a 405 nm diode laser with 30cycles of 1s 295ms resulting in a light dose of 6 J/cm^-2^. After irradiation the cells were imaged for 24 h with the Olympus IX81 inverted microscope (Olympus Life Science) using a 20x/0.75 NA air objective with an ORCA-ER CCD digital camera (Hamamatsu Photonics, Japan). Transmission, mCitrine and mCherry fluorescence images were acquired every 15 minutes.

In transgenic bCtC-DD HeLa cells, mCitrine and mCherry intensities were normalized to the mean intensity value at 4 h after 405nm illumination of NvocTMP-Cl treated cells.

In HeLa cells transiently transfected with bCtCDD-Hal and TOPmCherry, used to check for transcriptional activity after 405 nm irradiation of NvocTMP-Cl, cells were handled as above and imaged every 15 minutes. Fluorescence intensities were measured at every hour (1,2,3,4 h) for cells and timepoints without apparent cell division. Fluorescence intensity was Min-Max normalized in the 0-100% range between the smallest and the largest value in the dataset.

### Mouse small intestinal organoid (mSIO) culture

Wildtype mSIOs were generated and maintained as described by Sato et al., 2009 via isolation of crypts from a five month old C57BL/6N female mouse and maintained in 60% Matrigel (BD Corning) droplets covered with Advanced DMEM/F12 with 2 mM Glutamax, 10 mM HEPES, 100 units/mL Penicillin/Streptomycin (AdDF+++) with 1.25 mM *N*-acetyl-L-cystein, 1x B27, 50 ng/mL murine EGF, 100 ng/mL recombinant murine Noggin (Peprotech, # 250-38) and 500 ng/mL recombinant murine R-spondin 1 (Peprotech, # 315-32), this media is further named ENR. An additional batch of wildtype mSIOs, for local activation experiments, was purchased from STEMCELL Technologies (#70931; Cologne, Germany). The mSIOs were subcultured by mechanical disintegration every 4-5 days. To generate stably transfected transgenic mSIOs, PiggyBac-backbone based constructs were co-transfected together with the PiggyBac transposase in a 3:1 ratio of gene-of-interest construct to transposase by magnetofection via Lipofectamin 2000 (Thermo Fisher Scientific) and CombiMag^TM^ (BIOZOL Diagnostica). Pre transfection treatment: Four days prior transfection mSIOs were sub-cultured and after two days of growth treated with GSK3-inhibitor (CHIR99021 6 µM, Sigma-Aldrich) for 2 days. Post transfection treatment: GSK3-inhibitor (CHIR99021 6 µM, Sigma-Aldrich) and 10 µM RhoK inhibitor (Y-27632; Hölzel biotech) in ENR for 48 h. Alternatively, transfection was performed via electroporation (NEPA21 Electroporator; NEPA GENE) analog to the protocol by Fujii *et al* (2015). 48 h after transfection, mSIOs were selected for genomic integration by culture in presence of puromycin (1-2 µg/mL, Sigma-Aldrich). Cryopreservation of organoid lines was accomplished by freezing non-distintegrated organoids in Recovery™ Cell Culture Freezing Medium (Gibco™, (Thermo Fisher Scientific). Optional: addition of 10 µM RhoK inhibitor (Y-27632; Hölzel biotech) to the Freezing Medium.

Transgenic organoids harboring the pPB-bCtC-DD contruct are also abbreviated as ‘trans’ in some figures and text. Transgenic organoids with the pPB-bCtC are also referred as bCtC mSIOs.

### Preparation of Wnt3a-conditioned media

The Wnt3a producing cell line L-Wnt3a was gift from the Clevers lab (Hubrecht Institute / KNAW). The Wnt3a conditioned medium was produced as previously described (Pleguezuelos-Manzano *et al*, 2020) and mixed to: 50% Wnt3a-CM and 50% AdDF+++ with 1.25 mM *N*-acetyl-L-cystein, 1x B27, 50 ng/mL murine EGF, 100 ng/mL recombinant murine Noggin (Peprotech, # 250-38) and 500 ng/mL recombinant murine R-spondin 1 (Peprotech, # 315-32), further mentioned as 50% Wnt3a-CM.

#### Global, chemical βCtn4A-DD stabilization in mSIOs

In case of time-lapse experiments for early, acute chemical stabilization of βCtn4A-DD, fully developed bCtC-DD mSIOs were mechanically disintegrated, as described, and passed through a 70 μm cell strainer to exclude large organoid fragments. Wt and bCtC mSIOs were handled analogously. Resulting fragments are called crypt precursors. In case of late exposure experiments, the mSIOs were not passed through a cell strainer to enrich the diversity of developmental states at 48 h. The crypt precursors were seeded into 60% Matrigel droplets of 30-50 µl into glass-bottom plates or multiwell-slides and cultured in ENR.

In early, chemical βCtn4A-DD stabilization, media was changed after 3 h to Vehicle (IF-experiments: 0.1% DMSO in ENR; live cell imaging experiments: ENR), TMP (10 µM; solvent of 10mM TMP-stock solution in IF-experiments: DMSO; solvent of 1mM TMP stock solution in live cell imaging experiments: PBS) or Wnt3a (50% Wnt3a-CM). Media was refreshed every 48 h and mSIOs cultured for up to 120 h. In case of transient administration of Wnt3a (Wnt[48,-72]), the removal was performed by three washes with fresh ENR with 30 min incubation/wash. In case of transient TMP (TMP[48,-72]), the washout was assisted by 25 µM ecDHFR in ENR. After the washes, organoids were further cultured in ENR. In experiments with late exposure, ENR Media was changed to TMP (10 µM) or 50 % Wnt3a-CM at 48 h of mSIO culture.

### Live imaging of mSIOs after global, chemical βCtn4A-DD stabilization

Timelapse live imaging of mSIOs was performed on the Olympus IX81 inverted microscope (Olympus Life Science) equipped with an MT20 illumination system and an incubation chamber at 37 °C and 5% CO_2_ atmosphere. Using a 10x/0.4 NA or 20x/0.75 NA air objective and an ORCA-ER CCD digital camera (Hamamatsu Photonics, Japan). z-stacks of 120 µm with 6 slices were acquired in Transmission, mCitrine (YFP filter set) and mCherry (RFP filter set) every 1-2 h for up to 120 h organoid lifetime.

### Local, opto-chemical βCtn4A-DD stabilization in mSIOs

bCtC-DD transgenic organoids were treated with 10 µM NvocTMP-Cl or DMSO as Vehicle control for 90 min and then washed 3x by cycles of 1.) 3x 5 min incubation with fresh AdDF+++ media followed by 2.) 30 min incubation in ENR media to remove unbound compound. Resulting in a washing time of 2.25 h in total.

Photo-uncaging of NvocTMP-Cl was performed at an in-house combined setup consisting of an Olympus IX81 inverse microscope (Olympus Life Science), pE-4000 LED illumination system (CoolLED), Mosaic3 Digital Mirror Device (Andor), Stage Top Incubator (ibidi) and ORCA-Quest qCMOS camera (Hamamatsu Photonics, Japan). The device was controlled by μManager 1.4 Software (Edelstein *et al*, 2014). Illumination dose = (2.4mW x 4s) / 88657 µm^2^ = 10.8 J/cm². Uncaging of the compound in selected organoid regions was followed by 48-72 h of imaging using a 20x/0.75 NA air objective. Transmission, mCitrine and mCherry fluorescence image z-stacks were acquired with 1 h/frame.

### Immunofluorescence

For fixation, organoids were washed 5x with 1x PBS (phosphate buffered saline) at RT before fixation with 4% PFA (paraformaldehyde) for 15 min at RT. In case of remaining Matrigel, 4% PFA treatment can be repeated up to two more times for max 45 min in total. After 3x washes with 1x PBS, permeabilization was performed using 0.5% Triton X-100 in PBS for 1 h at RT, followed by blocking with 3% BSA (bovine serum albumin) with 0.1% Triton X-100 in 1x PBS for 1 h at RT. Primary Antibodies: rabbit anti-Lysozyme (#A0099, Dako), rabbit anti-AldolaseB (#ab75751, Abcam), mouse anti-β-catenin mAb (sc-7963, Santa Cruz Biotechnology). Secondary Antibodies: donkey anti-mouse-AlexaFluor-488 or -546 pAb (Invitrogen), donkey anti-rabbit-AlexaFluor-546 pAb (Invitrogen). F-Actin Staining: Phalloidin-AlexaFluor-647 (Invitrogen); Nuclei: DAPI (20 μg/mL). Fructose–glycerol clearing solution: 60% (vol/vol) glycerol and 2.5 M fructose. Refractive index = 1.4688 at RT (Dekkers *et al*, 2019).

### Confocal laser scanning microscopy (CLSM)

mSIOs were cultured, fixed and stained by immunofluorescence as described using 8-well Lab-Tek™ chambered cover glass slides (ThermoFisher). Confocal image stacks were acquired using a Leica TSC SP8 microscope (Leica Microsystems, Wetzlar, Germany), which was equipped with a 405-nm diode laser and a white light laser (white light laser Kit WLL2, NKT Photonics). Objective: HC PL APO CS2 20x/0.75 DRY; Pinhole: 1.0 Airy units. Excitation wavelengths: DAPI (405nm), AlexaFluor 488 (488nm), Alexa Fluor 546 (560nm), Alexa Fluor 647 (640nm). Fluorescence emission was detected by hybrid detectors (HyD) restricted to: DAPI(410-463), AlexaFluor 488 (506-552nm), Alexa Fluor 546 (568-615nm), Alexa Fluor 647 (658–720 nm). With 200-400 Hz scanning frequency, 12-bit images of 512 × 512 pixels were recorded.

### Western blot analysis

Cell and organoid lysates were prepared by lysing sample pellets in M-PER lysis buffer (Thermo Fisher Scientific) supplemented with phosphatase inhibitor cocktail 2 and 3 (P5726, P0044; Sigma Aldrich) and cOmplete™ mini protease inhibitor cocktail (Roche). Protein concentrations were measured with the Micro BCA Protein Assay (Thermo Fisher Scientific) against a bovine serum albumin standard curve for equal loading of protein samples. Proteins were resolved by SDS PAGE using 10% PAA gels and transferred to Immobilon P PVDF membranes (Millipore) or nitrocellulose 0.45 µm (Thermo Scientific) membranes using the Invitrogen Mini Gel Tank and Blot Module (Invitrogen). Protein load: 30 μg. Tris-Glycine-SDS Running buffer: 0.1% SDS, 25 mM Tris, 0.19 M Glycine. Towbin Transferbuffer: 25 mM Tris, 0.19 M Glycine, pH 8.6, 18% Methanol, 0.025% SDS. Blocking buffer: 5% BSA in 50% LI-COR intercept TBS-T blocking buffer (LI-COR Bioscience), 50% TBS-T. Primary antibody incubation overnight at 4°C, secondary antibody incubation 45 min at RT. Antibodies: goat anti beta-Catenin pAb (R&D Systems, AF1329; 1:1000 dilution), rabbit anti Axin2 mAb (Cell Signaling; 2151; 1:1000 dilution), rabbit anti α-Tubulin (Sigma Aldrich, T6074; 1:10,000 dilution). Secondary antibodies: IRDyes® 680RD or 800CW Donkey anti-Goat IgG (H + L), Donkey anti-Rabbit IgG (H + L). Blots were scanned with the Odyssey Infrared Imaging System (LI-COR Bioscience). Western blot images were analyzed using LI-COR Image Studio Software.

Quantification of Western Blot

Mean integrated intensity of protein bands was used for analysis. Protein bands were normalized by mean intensity over all samples on the membrane and relative to Tubulin as reference protein. POI: Protein of interest

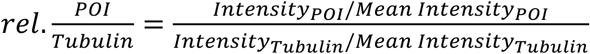

### Image Analysis

#### Paneth Cell Positions and Distance Matrices (DMs)

Paneth Cell (PC) positions were extracted from confocal micrograph z-stacks of mSIOs stained for lysozyme by using a self-developed macro in FIJI (Version 1.53a-1.54p) (Schindelin *et al*, 2012). The main steps of the macro are: 1. Maximum intensity projections (MIPs) of lysozyme were used to extract PC positions in xy (µm scale) by “Find Maxima” command with threshold adapted to the experiment’s signal. 2. The corresponding z-position was determined by determination of the maximum intensity in z for each xy PC position. PC positions in z were recalculated from pixel to µm scale by using image voxel size.

To calculate the corresponding distance matrix the PC positions were first sorted by proximity. Next, the Euclidean distances of each PC to every other PC were calculated using matlab’s pdist function and converted into a squared form where the diagonal of zeros represents the distance of each PC to itself (Fig. EV3D). Dark areas close to the DM diagonal indicate clusters of PCs while areas of lighter shades indicate the relation of the PC clusters to each other.

These resulting full distance matrices (DMs), excluding the diagonal of zeros, were used to calculate statistics on the PC distance distribution like the normalized standard deviation (nSD) (SD of DM/Mean of DM) and kurtosis (KT). Shannon entropy (SE) is used as a variable representing regularity in PC cluster arrangement. Full DMs (32-bit) were binarized by first applying FIJIs “Find Edges” command, this highlights the transitions between areas of similar distances, followed by application of an auto-threshold (Li; Li’s Minimum Cross Entropy thresholding) to generate a binary representation of PC clusters. Minimal full DM size was set to 15 PCs, Organoids and DMs with less PCs were excluded to prevent bias by underdeveloped mSIOs and enteroids. Shannon Entropy (SE) of these binary DMs serves as a parameter to assess regularity of the PC clusters (Fig. EV3E) with low values indicating regular distribution of clusters and high values an elevated irregularity.

### Average Phase-matched Distance Matrices (APDMs)

In prototypically developed mSIOs with crypt-villus structure, a range of 30 PCs usually encompasses 2-3 crypts. Per population, separated by genetic background and treatment, sub-matrices of 30x30 PCs along the diagonal were matched by minimizing the difference within the population. For this, the smallest prototypical DM/condition was used as a template to match the most similar sub-matrix of every DM. By this, each full DM generated a 30x30 sub-matrix, population sub-matrices were averaged to generate the APDM. APDMs represent a common PC-organization within a mSIO population.

### Morphometric Analysis mSIO image segmentation

Segmentation of confocal micrographs from mSIOs stained by immunofluorescence was performed in Fiji with self-developed macros. The main steps are: Confocal z-stacks were projected by maximum intensity (MIP), creating a composite image of all channels. The MIP composite was flattened, projecting all channels to an RGB image. This RGB image was converted to 8-bit, followed by auto threshold method “Percentile dark” to create a binary mask. The resulting contours were smoothed by interpolation (10px), shown as overlay on the initial projection and corrected for crypt separation and removal of debris.

Segmentation of widefield micrographs from mSIOs in live-imaging experiments was performed in Fiji with self-developed macros. The main steps: brightfield z-stacks were projected by median intensity, edges detected via “Find Edges” command and converted to binary via "Triangle dark" auto-thresholding. These raw masks were further converted via “Close” (iteration of 3) and “Fill Holes”. These masks were again smoothed and corrected as described above.

### Morphometric feature extraction

For morphometric classification 32 scale- and rotational independent features were extracted from the binary mSIO masks.

### FIJI Shape descriptors (4)

Circularity: 4π*area/perimeter^2^; Aspect Ratio: major_axis/minor_axis; Roundness: 4*area/(π*major_axis^2^); Solidity: area/convex area.(Schindelin *et al*., 2012)

#### Hu-Image Moment Invariants 1-7 (7)

(Hu, 1962; Huang & Leng, 2010), calculated from the central image moments acquired by the Fiji Image Moments plugin from Dimiter Prodanov (2008).

**Zernicke Moments** (Teague, 1980)**(16)** Z_n_^-m^ for n=0-6, m ≤ n (16 moments) with a Matlab Script by Amir Tahmasbi (Saki *et al*, 2013; Tahmasbi *et al*, 2011)(2016). Z_n_^m^: Z_0_^-0^; Z_1_^1^; Z_2_^0^; Z_2_^2^; Z_3_^1^; Z_3_^3^; Z_4_^0^; Z_4_^2^; Z_4_^4^; Z_5_^1^; Z_5_^3^; Z_5_^5^; Z_6_^0^; Z_6_^2^; Z_6_^4^; Z_6_^6^;

To generate the combined feature “**Number of crypt-like features / Area fraction of largest inscribed circle**” **(1)** the ratio of the two features “**Number of crypt-like features**” and “**Area fraction of largest inscribed circle**” was calculated.

#### Area fraction of largest inscribed circle

The largest inscribed circle was fit into the segmented binary organoid by using the MorphoLibJ plugin (Legland *et al*, 2016). The Area fraction was calculated by Area_Circle_/Area_Organoid_ and serves as an estimator of the relative villus size.

#### Number of crypt-like features

The organoid contour was linearized by measurement of the distance between the contour and the largest inscribed circle on the radius of the organoid contour. This distance was plot against the organoid contour length (perimeter). Crypts and crypt-like features presented as peaks in this plot, which could be detected via the findpeaks command in Matlab. Specifying parameters: minimal peak prominence 15μm; maximal peak width 1/3 of perimeter.

#### Relative Peak Width/Height (2)

Peak Width and Peak Prominence represent dimensions of the crypts and crypt-like features. Both were extracted from the detected peaks of the linearized contour above in Matlab. Relative Peak Width was calculated as ratio of Peak Width to the Perimeter. Relative Peak Height was calculated as ratio of Peak Prominence to the maximal axis length (width or height).

#### Convex/concave fraction of contour (2)

Convexity/concavity of contour was calculated by using the determinant (det) for three adjacent points along the full contour. For det>0 that region of the contour was concave, for det<0 convex and for det=0 collinear.

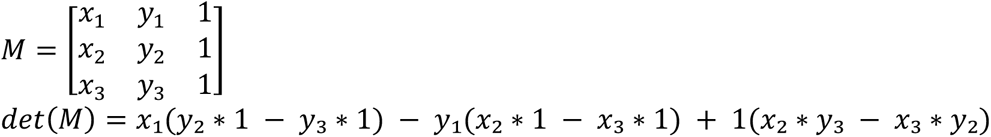

### Reduction of morphometric feature set

The 32 morphometric features were reduced by removal of highly correlating features and elimination of features with low predictive power as determined by sequential feature selection using different classifier models. For example, from Circularity, Solitidy, Roundness and Aspect Ratio only the Aspect Ratio with a high predictive power for organoid morphologies was selected. That resulted in the following 8 features to generate a morphometric feature space suitable to differentiate organoid morphologies: Number of Crypt-like Features / Area-fraction of Largest Inscribed Circle, Relative Peak Width, Aspect Ratio, Hu Moment 1, Hu Moment 2, Hu Moment 4, Zernicke Moments Z_2_^0^ and Z_2_^2^.

### Principal Component Analysis (PCA) of the morphometric feature set data

PCA as a linear data transformation is used to visualize distribution of endpoint morphologies in an optimal morphometric space. The PCA was performed by Matlabs pca function, using the 8 morphometric feature set of the mSIOs stained by indirect immunofluorescence. The resulting coefficients/loadings were subsequently applied to transform the morphometric data from other experiments into the same principal component space. The optimal three dimensional morphometric space is created by the three components PC1, PC2, PC3.

### Morphometric ground truth generation

The morphometric ground truth was generated by transformation of population/treatment-related mSIO clusters from the Paneth Cell organization space of Shannon entropy (SE), as marker of PC cluster regularity, against two statistical moments of PC distances (nSD: normalized SD, KT: kurtosis) (Fig. EV3F) into the morphometric space (PC1, PC2, PC3; Fig. EV3 I). First, treatment groups of mSIOs were combined if they were not significantly different for the three parameters (nSD, KT, SE). By this ‘controls’ is a combination of wt Vehicle, wt TMP and trans Vehicle, ‘Wnt’ is the combined mSIOs of wt Wnt and trans Wnt, while trans TMP and bCtC formed their own treatment groups. From these the core fraction of mSIOs was selected by using the mSIOs within mean±2xSD volume of all three PC-organization parameters (nSD, KT, SE; Fig. EV3 F). This core fraction was used to train a k nearest neighbour (knn)-based classifier which was subsequently applied to classify all mSIOs (Fig. EV3 G). The confusion table shows, that the treatment groups partially share features of PC organization and no group is completely distinct from the others. mSIOs from the diagonal of the resulting confusion matrix were classified according to their treatment and such considered as representative PC-organization for this treatment group. Projected into the morphometric principal component space (PC1, PC2, PC3) the ‘controls’ provided the morphometric volume for the prototypical crypt-villus morphology of normally developed mSIOs, further referred to as ‘Regular’ morphology. The morphometric volume covered by the bCtC mSIOs was considered as the ‘Cystic’ region, due to the round, cryptless appearance. The class labels were further refined by keeping only mSIOs which were inside the corresponding mean±SD region for at least two dimensions of PC1, PC2 and PC3. The removed mSIOs were considered as unknown morphology and combined with the projected representative mSIOs from the ‘Wnt’ and ‘trans TMP’ populations. A k-means clustering (De Landtsheer, 2019) was applied to this data, including determination of optimal cluster number by elbow method, to identify a clustering of the morphometric groups outside of the ‘Regular’ and ‘Cystic’ regions. This resulted in three additional morphometric clusters (Fig. EV3 I) which harboured mSIOs with the following features: absence of crypts, enriched in crypt-like structures or enlarged crypt-like structures. In total 97 out of 170 mSIOs were considered as representative regarding PC-organization for their treatment and were part of one of 5 clusters in the morphometric space building the ground truth of morphologies.

### Classifier Training and Testing

Based on the identified ground truth of mSIO morphologies, using the Matlab classification learner app, different simple classifiers were trained with the 8 morphometric features as predictors and rated by accuracy to decide for the most robust classifier structure. The final supervised Classifier Training was performed with an 8-fold cross validation on an optimizable Decision Tree classifier. The cross-correlation table of this classifier and its statistical parameters are shown in Fig. EV3 J. For Classification this classifier is applied on the 8-dimensional morphometric feature set of mSIOs.

### Fluorescence Analysis of βCtn4A-DD stabilization in mSIOs

The oncoprotein βCtn4A is fused to the mCitrine fluorophore in βCtn4A-DD. The fluorescence intensity can thus be used as approximation for the amount of stabilized protein. mSIO development was imaged as described above on an Olympus IX81 inverted widefield microscope. From the acquired z-stack, maximum intensity projections were generated. Segmented binary mSIO masks were used to measure the corresponding mean intensity in the peripheral cell layer by shrinking the mSIO contour by 10px and converting the area selection to a line selection with line thickness of 10px. This line selection coveres the peripheral cell layer. Since βCtn4A-DD protein is also stabilized in the cell-cell contacts of the villus region, βCtn4A-DD intensity in TMP or NvocTMP-Cl treated mSIOs was related to the mean intensity of the corresponding Vehicle (DMSO) controls.

### Fluorescence Intensity upon local βCtn4A-DD stabilization

Local βCtn4A-DD fluorescence intensity after NvocTMP-Cl incubation and irradiation with 405 nm was measured on projections of z-stacks acquired on an Olympus IX81 inverse widefield microscope, as described above. Since the goal of this analysis is not quantitative but to follow the flow of fluorescent material by local intensity changes over the organoid development, images were preprocessed to correct for the illumination profile and normalize the values. This preprocessing was performed by application of a gaussian blur with a radius of 500 px (pixel size: 0.156 µm x 0.156 µm) to generate the illumination profile. Each Channel (mCitrine, mCherry) was divided by this illumination profile to generate the corrected z-stacks. On these corrected z-stack, we applied an average z-projection and measured the local intensities in the peripheral cells on the shrunken contour (-35px) as line selection (10px width) as described above. To correct for growth and match corresponding regions, crypts and villus regions were separated. To correct for the growth, these separated regions were resampled to the size at 18 h after 405 nm irradiation (total 66 h). The resulting intensity around the organoid periphery was sorted to center the irradiated precursor/crypt region within the contour, smoothed with a moving average span of 100px and background corrected by subtraction of the minimum value per timepoint. Per organoid, this size normalized fluorescence was plotted and additionally summed up to create the cumulative fluorescence curve. The cumulative fluorescence was corrected to the value of postPA PAmCherry, since that signal is only created at the timepoint of PA and further distributed and diluted by cell division. Thus, the cumulative fluorescence at each timepoint of the βCtn4A-DD signal was corrected by the factor of the corresponding PAmCherry/ postPA PAmCherry.

### Statistics

Statistical tests were calculated via GraphPad Prism version 10 for macOS, GraphPad Prism 10 for macOS (v10.6.1), Boston, Massachusetts USA, www.graphpad.com. Tests are indicated in the corresponding figure legends.

## Acknowledgements

This project was supported by Max Planck Society for the Advancement of Science. We would like to thank Manuela Grygier and Jutta Luig for their assistance in cloning and plasmid preparation; Xi Chen and Yaowen Wu for the critical assistance in method development; Roger Goody, Jan Hübinger, Malte Schmick, Mario Caracci Oporto and Luis Munoz Nava for critically reading the manuscript, feedback and discussions. Sven Müller for technical support with confocal microscopy and photoactivation setup. Prof. Dr. Philippe I.H. Bastiaens, passed away on May 15, 2025. All authors agree that his inclusion as a co-author is appropriate to honor his intellectual contribution to this work.

## Disclosure and competing interests statement

No competing interests to declare.

## Funding

Project ’Oncomeme’ (2018-2021, Max Planck Society -funds for special needs)

## Author contributions

Birga Soetje: Conceptualization (evolution); methodology; software; formal analysis; investigation; visualization; data curation; writing – original draft; writing – review & editing.

Hui Ma: Conceptualization (original); methodology (creation of models); formal analysis; investigation; writing – review & editing.

Sarah Imtiaz: Conceptualization (evolution), methodology; data curation; formal analysis; investigation; visualization, writing – review & editing.

Agustin A Corbat: Investigation; formal analysis; methodology; software; writing – review & editing.

Hernan E Grecco: Funding acquisition (of AAC); methodology; software; supervision (of AAC); visualization.

Lisaweta Schröder: Formal analysis; investigation; visualization. Yannick Brüggemann: Investigation.

Sabrina Seidler: Investigation. Michael Reichl: Investigation.

Philippe IH Bastiaens: Conceptualization (original and evolution); supervision; methodology; formal analysis; investigation; visualization; writing – original draft; project administration; funding acquisition.

## Data Availability

Self-developed macros and matlab scripts will be provided on github.

## Supplementary information

Video1: bCtC organoid in Transmission, 24-108h.

Video2: bCtC-DD organoid in Transmission, bCtn4A-Cit and PAmCherry after cryptPA, 48-99h.

Appendix Figures S5&S6:

**Appendix Fig. S5. Stages of mSIO development and visualization of βCtn4A-DD stabilization.**

**Appendix Fig. S6. Scatterplots of cumulative βCtn4A-DD fluorescence versus crypt perimeter.**

---

**Figure EV2.**
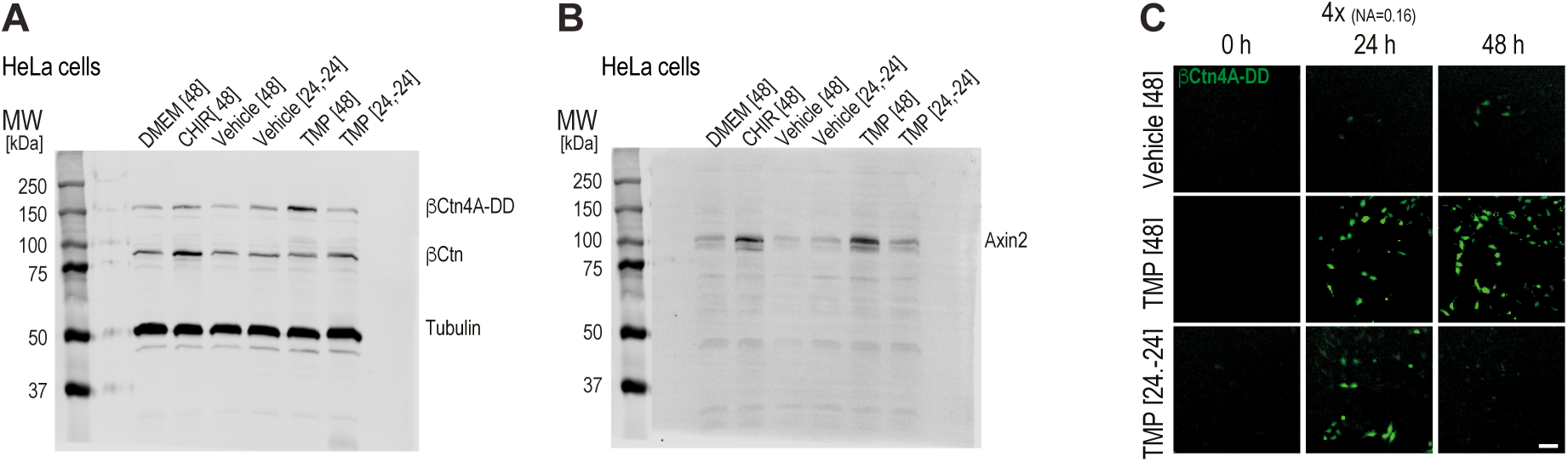
Bioorthogonal chemical control of oncogenic β-Catenin in HeLa cells. **A.** Representative western blot for data in Fig.2A: βCtn4A-DD and endogenous β-Catenin (βCtn) in transgenic HeLa bCtC-DD and tubulin as reference protein (700 nm channel). DMEM[48]: 48h full growth media; CHIR[48]: 48 h DMEM + 5 µM CHIR99021; Vehicle[48]: 48 h DMEM+0.1% DMSO; Vehicle[24,-24]: 24 h DMEM + 0.1% DMSO followed by washout and 24 h DMEM; TMP[48]: 48h DMEM + 10 µM TMP; TMP[24,-24]: 24 h DMEM + 10 µM TMP followed by washout and 24 h growth in DMEM. MW[kDa]: molecular weight makers in kilo Dalton **B.** Representative western blot as in (A) for Axin2 (800 nm channel) for data in Fig.2B. **C.** Representative micrographs of βCtn4A-DD fluorescence at indicated time points in transgenic bCtC-DD HeLa cells for Vehicle[48], TMP[48], TMP[24,-24]. scale bar: 100 µm.

**Fig. EV3.**
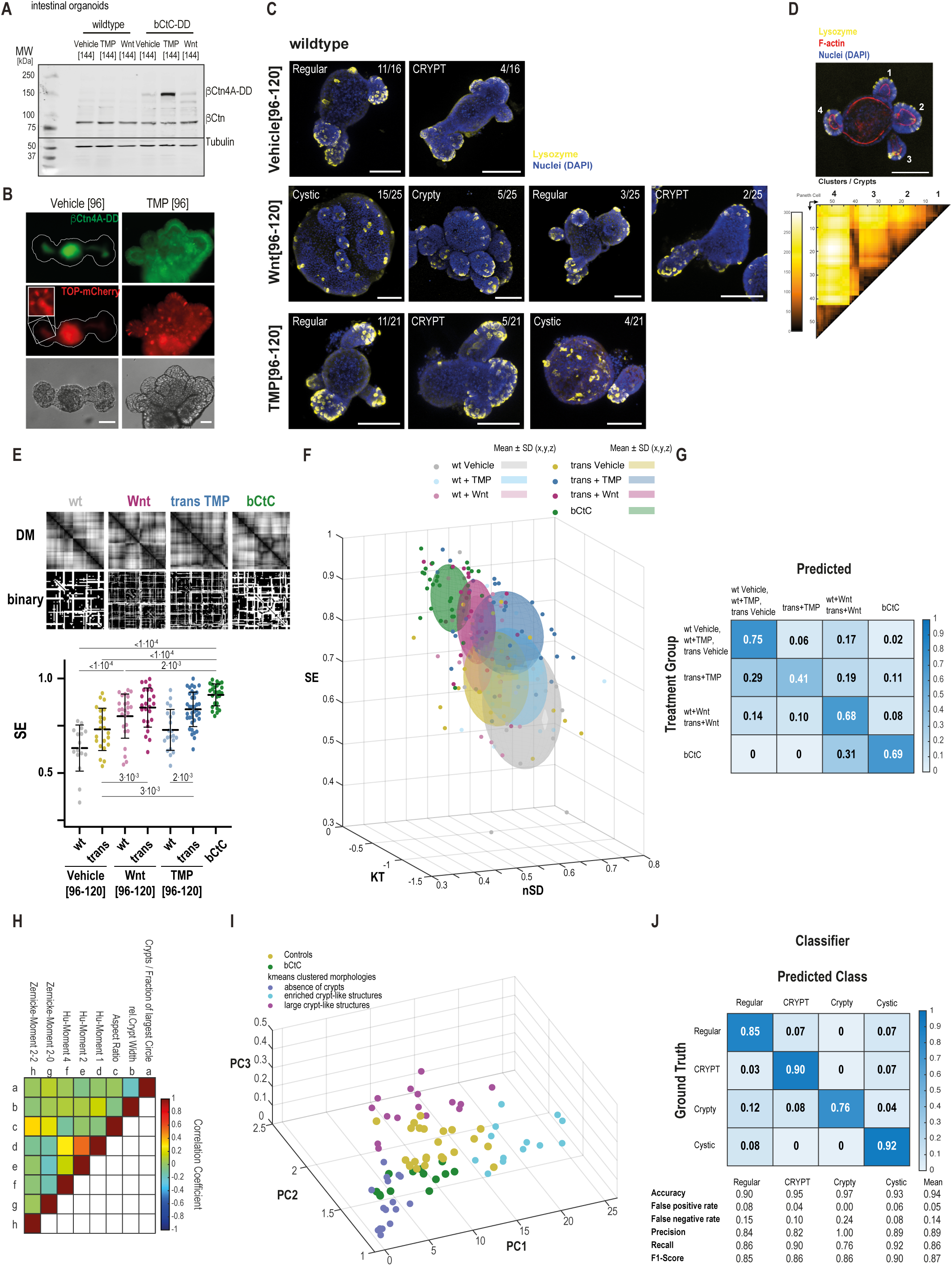
Mapping Paneth cell organization in mSIOs to morphometric classes. **A.** Corresponding western blot to Fig. 3A showing βCtn4A-DD and endogenous β-Catenin (βCtn) in wildtype (wt) and transgenic bCtC-DD mSIOs. Tubulin(inset): reference protein. Vehicle[144]: 144 h ENR + 0.1% DMSO; TMP[144]: 144 h ENR + 10 µM TMP, Wnt[144]: 144 h ENR + 50% Wnt3a-conditioned media. **B.** Representative micrographs of βCtn4A-DD (top, green) or TOP-mCherry (center, red) transcriptional reporter fluorescence in Vehicle[96] or TMP[96] bCtC-DD-Hal+TOP-mCherry small intestinal organoids. TOP-mCherry inset: intensity enhanced zoom of the crypt. Bottom: transmission images. Scale bar: 100 µm. **C.** Representative maximum intensity projections (MIPs) of 3D CLSM immunofluorescence stacks of wildtype mSIOs. Yellow: Paneth cell marker antibody α-lysozyme, Blue: Nucleus marker DAPI. Rows from top to bottom: Vehicle[96-120]: 96-120 h ENR + 0.1% DMSO; Wnt[96-120]: 96-120 h ENR + 50% Wnt3a-conditioned media; TMP[96-120]: 96-120 h ENR + 10 µM TMP. N=4/3/4; n=16(2/4/2/8)/25(6/3/16)/21(1/1/2/17). Mclasses and corresponding fractions indicated at top of MIPs in white. Scale bar: 100 µm. **D.** Maximum intensity projection (MIP) of 3D CLSM immunofluorescence stacks of wildtype mSIO. Yellow: Paneth cell marker antibody α-lysozyme, Blue: Nucleus marker DAPI. Red: F-Actin by Phalloidin. scale bar: 100 µm. Corresponding distance matrix (DM) of Paneth cells. Color bar: PC distance in µm. Numbered structures in the micrographs correspond to numbered areas in the DMs. **E.** Top: DMs and corresponding edge-detected, binary DMs exemplary for the median of the four β-catenin activity groups in wildtype (wt) and transgenic bCtC-DD (trans) mSIOs. wt: wt Vehicle[96-120], wt+TMP[96-120], trans Vehicle[96-120]; Wnt: wt + Wnt[96-120], trans + Wnt[96-120]; trans TMP: trans + TMP[96-120]; bCtC: bCtC mSIOs. Bottom: Shannon Entropy (SE) distribution of edge-detected, binary DMs from wt and bCtC-DD (trans) in Vehicle[96-120], Wnt[96-120], or TMP[96-120], and bCtC[96] mSIOs. p-value: one-way ANOVA; Tukey’s multiple comparisons test **F.** 3D scatter plot of DM’s statistical parameters from wt and bCtC-DD in Vehicle[96-120], Wnt[96-120], TMP[96-120], and bCtC mSIOs. x-axis: normalized Standard deviation (nSD; SD/Mean); y-axis: Kurtosis (KT); z-axis: Shannon Entropy (SE). Ellipsoids of Mean±SD (x,y,z) **G.** Confusion matrix of KNN-based classifier trained on the points within Mean±SD(x,y,z) range of the 3D plot in (F). **H.** Correlation matrix of the reduced 8-dimensional morphometric feature-set applied for Mclassification. Color bar: correlation coefficient. **I.** 3D Principal component projection (PC1, PC2, PC3) of 8-dimensional morphometric parameter distribution for morphological ground truth determination. mSIOs at the diagonal in (G) of wt controls (wt Vehicle[96-120], wt+TMP[96-120], trans Vehicle[96-120]; yellow) and bCtC (green) serve as benchmark for two distinct PC-organization related morphological classes. Points outside of these two defined regions were clustered by k-nearest neighbors and cluster names assigned which reflect the prototypical phenotype of that cluster: absence of crypts (purple), enriched in crypt-like features (blue), large crypt-like features (magenta). **J.** Confusion matrix of a decision-tree based Mclassifier trained on the ground truth of four morphologies identified by PC organization and morphometric clustering in (I). bCtC and ‘absence of crypts’ clusters from (I) were combined to the ‘Cystic’ class during iterative classifier training. Mclassifier is trained using the 8 morphometric features in (H). Mclassification performance parameters accuracy, false positive and false negative rate, precision, recall and F1-Score calculated per Mclass and on average.

**Fig. EV4.**
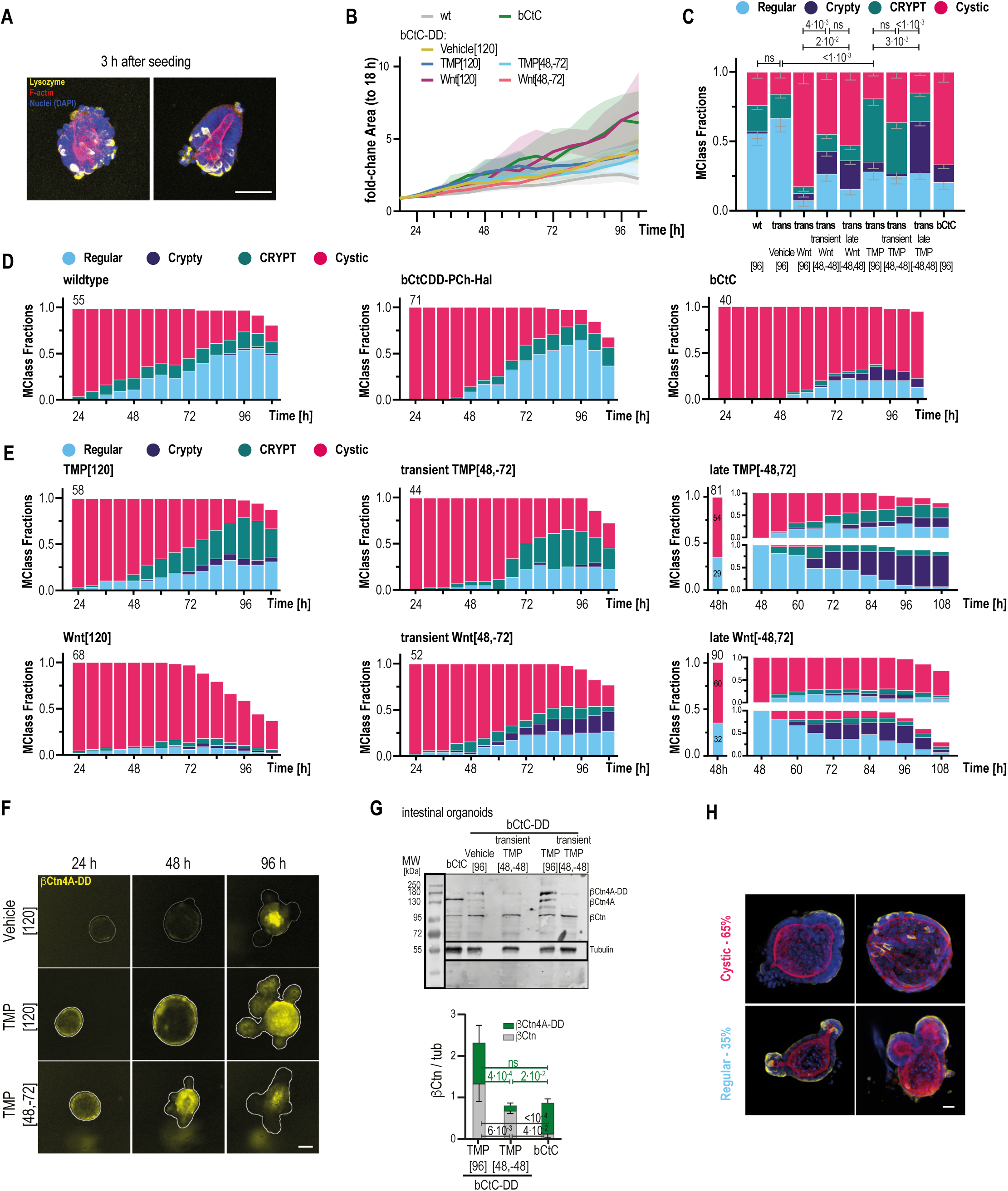
Temporal modulation of βCtn4A-DD stabilization and its morphological consequences. **A.** MIPs of immunofluorescent bCtC-DD organoid precursors 3 h after mechanical disintegration of fully developed mSIOs and seeding into Matrigel, representing the experimental initial state of organoid development. Lysozyme (yellow, Paneth Cells (PCs)), DAPI (blue, nuclei), F-Actin (red). Scale bar: 20 µm. **B.** Organoid growth as measured by fold-change to 24 h timepoint of cross-sectional area of organoid contours from segmented transmission images over time for wt, bCtC-DD Vehicle[120], TMP[120], transient TMP[48,-72] Wnt[120], transient Wnt[48,-72] and bCtC. Median ± 95% CI. N=4/4/4/4/3/3/4; n=55/52/71/58/42/39/40. **C.** Stacked bargraph of Mclass fraction distributions at 96 h of wt[96], bCtC-DD (trans) Vehicle[96], Wnt[96], transient Wnt[48,-48], late Wnt[-48,48], TMP[96], transient TMP[48,-48], late TMP[-48,48] and bCtC[96]. N=4/6/3/3/4/4/4/4/4; n=54/69/40/49/83/57/44/73/39. p-values: chi-squared test. **D.** Stacked bargraph of Mclass fraction distributions over time for every 6 h from 24 h to 108 h for wildtype (left; N=4; n= 55(10/8/18/19)), bCtC-DD Vehicle[120] (center; N=6; n= 71(15/14/19/15/3/5)) and bCtC (right; N=4; n=40(11/9/11/9)) mSIOs. **E.** Stacked bargraph of Mclass fraction distributions over time for every 6 h from 24 h to 108 h for bCtC-DD mSIOs exposed to TMP[120] (top left; N=4; n=58(20/23/8/7)), transient TMP[48,-72] (top center; N=4; n=44(15/20/3/6)), Wnt[120] (bottom left; N=3; n=68(27/22/19)), transient Wnt[48,-72] (bottom center; N=3; n=52(24/18/10)), or for every 6 h from 48 h to 108 h for late TMP[-48,72] (top right; N=4; n=81(21/18/19/23)), late Wnt[-48,72] (bottom right; N=4; n=90(16/22/21/31). **F.** Micrographs of βCtn4A-DD in developing bCtC-DD organoids at indicated time points and treatment: Vehicle[120], TMP[120], TMP[48,-72]. Scale bar: 100 µm. **G.** Western Blot: βCtn4A-DD and endogenous β-Catenin (bCtn) with Tubulin (inset) for reference in transgenic bCtC or bCtC-DD mSIOs for Vehicle[96], TMP[96] or transient TMP[48,-48]. Graph: Corresponding relative protein levels (βCtn/tub) βCtn4A-DD /Tubulin (green) and endogenous β-Catenin βCtn/Tubulin (grey). Mean±SD, N=4/5/3 per condition, p-values (color coded): two-way ANOVA, Sidak’s multiple comparison test. **H.** Representative maximum intensity projections (MIPs) of 3D CLSM immunofluorescence stacks of bCtC-DD mSIOs at 48 h with corresponding Mclasses representing the experimental initial state in late induction of oncoprotein activity by TMP[-48,72] or Wnt[-48,72]. Lysozyme (yellow, Paneth Cells (PCs)), DAPI (blue, nuclei), F-Actin (red). Percentage of mSIOs per Mclass ‘Cystic’ or ‘Regular’ indicated.

**Fig. EV5.**
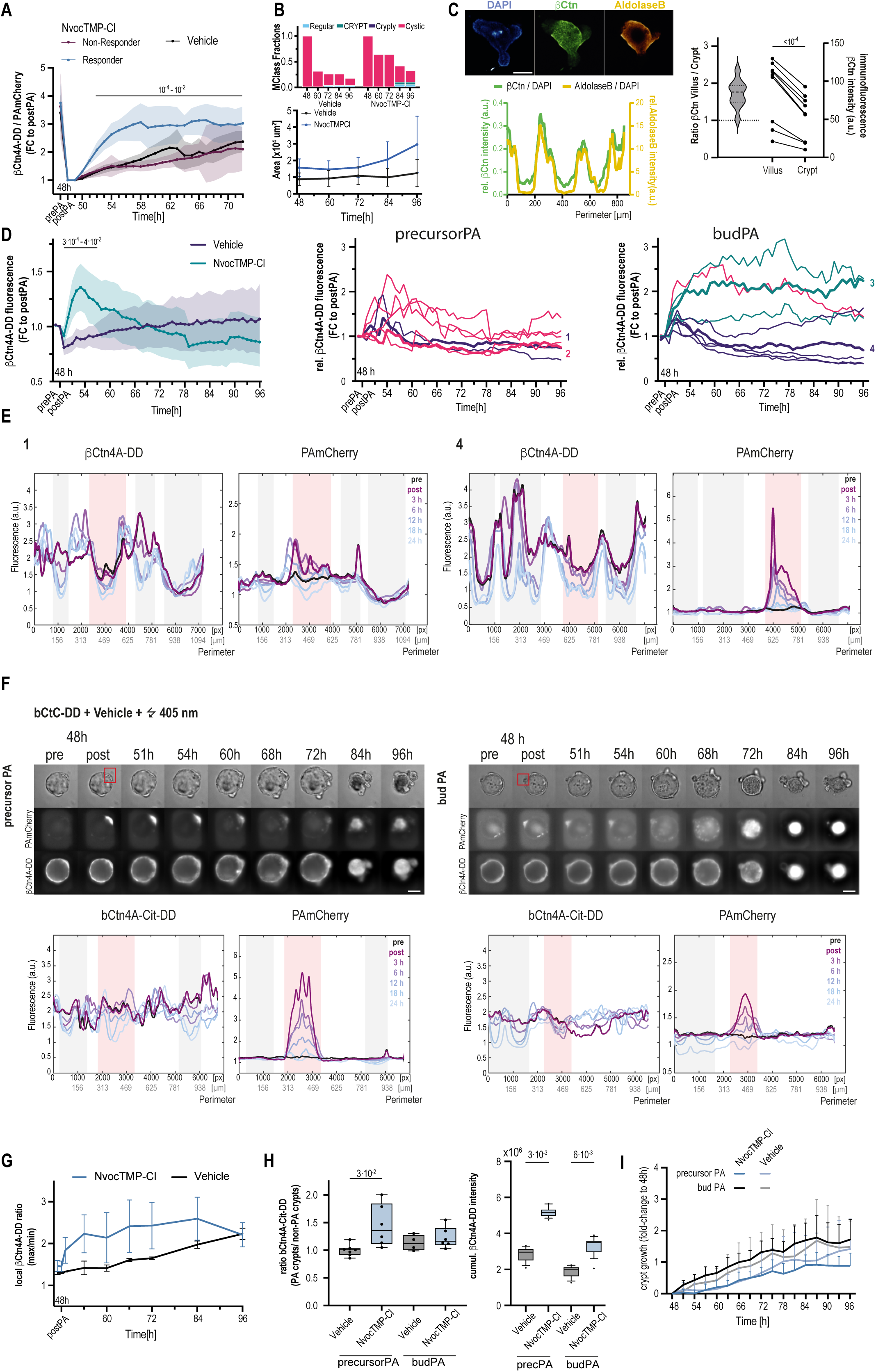
Consequences of opto-chemical oncoprotein stabilization in regenerating mSIOs. **A.** Expression ratio of βCtn4A-DD to PAmCherry of NvocTMP-Cl (10 µM) and Vehicle treated, entirely 405 nm irradiated (10 J/cm^2^) bCtC-DD mSIOs over time normalized to the postPA timepoint. NvocTMP-Cl treated mSIOs are separated by high (blue; Responder) and low (plum; Non-Responder) stabilization of βCtn4A-DD. Mean±95% CI. N=2; Nvoc-TMP-Cl n=15; Vehicle n=11. p-values: 2way ANOVA; uncorrected Fisher’s LSD of Responder against postPA timepoint. **B.** Mclass fractions (top) and organoid growth as crosssectional area over time (bottom; Mean±95% CI) of NvocTMP-Cl (10 µM) and Vehicle treated and entirely 405 nm irradiated (4s; 10 J/cm^2^) bCtC-DD mSIOs. **C.** Top: Maximum intensity projections (MIPs, top) and corresponding relative fluorescence along the contour (bottom) of 3D confocal laser scanning microscopy immunofluorescence stack of a bCtC-DD Vehicle [96] mSIO. Blue (DAPI): Nucleus marker DAPI; Green (βCtn): total β-catenin antibody α-β-catenin; Yellow (AldolaseB): Enterocyte marker antibody α-aldolase B. Bottom: Linearized relative β-catenin and aldolase B levels from exemplary MIPs along the contour showing difference in crypt and villus region expression levels. Right: Villus to crypt ratio distribution of β-catenin fluorescence (antibody α-β-catenin) in bCtC-DD Vehicle [96] mSIOs. N=3, n=10. p: Ratio-paired t-test. **D.** Left: Expression of βCtn4A-DD in NvocTMP-Cl (10 µM) and Vehicle treated and locally 405 nm irradiated (10 J/cm^2^) bCtC-DD mSIOs over time, normalized to the postPA timepoint. Mean±95% CI. N=5/5; n=16/16. p-values: 2way ANOVA; uncorrected Fisher’s LSD. Center, Right: Single mSIO βCtn4A-DD expression of NvocTMP-Cl (10 µM) treated relative to corresponding Vehicle-treated and locally, precursor PA (center) or bud PA (right), 405 nm irradiated (4s; 10 J/cm^2^) bCtC-DD mSIOs over time, normalized to the postPA timepoint. Color corresponds to end-stage Mclass: blue: ‘Regular’, violet: ‘Crypty’, green: ‘CRYPT’, red: ‘Cystic’. Thick lines with numbers correspond to mSIO contours and MIP sequences in Fig. 5 F,G. **E.** Quantification of local βCtn4A-DD and PAmCherry fluorescence along the developing mSIO contours for NvocTMP-Cl treated examples 1 (precursor PA) and 4 (bud PA) at indicated timepoints (prePA, postPA, 3 h, 6 h, 12 h, 18 h, 24 h) with normalized region sizes and matched regions (villus and crypt regions at 18 h after PA). Background color indicates regions: red – PA region; grey – additional crypts. **F.** Top: Micrographs of developing bCtC-DD Vehicle control mSIOs in Transmission, PAmCherry and βCtn4A-DD pre and post local irradiation (PA; red rectangle) at indicated timepoints for precursor PA (left; 1) and bud PA (right; 2) corresponding to the contours and trajectories in Fig. 5E. Bottom: Quantification of local βCtn4A-DD and PAmCherry fluorescence along the developing mSIO contours for the precursorPA and cryptPA at indicated timepoints (prePA, postPA, 3 h, 6 h, 12 h, 18 h, 24 h) with normalized region sizes and matched regions (villus and crypt regions at 18 h after PA). Background color indicates regions: red – PA region; grey – additional crypts. **G.** βCtn4A-DD fluorescence intensity ratio over time from local maxima to local minima of the exemplary (Fig. 5E,F) NvocTMP-Cl (10 µM; blue) and Vehicle (black) treated and locally 405 nm irradiated (4s; 10 J/cm^2^) bCtC-DD mSIOs. Mean±SD. NvocTMP-Cl : n=4; Vehicle: n=2. **H.** Left: βCtn4A-DD fluorescence intensity ratio of PA versus non-PA crypts of the same mSIO. Boxplot Min-Max. Precursor PA: Vehicle n=7; NvocTMP-Cl n=6; bud PA: Vehicle n=4; NvocTMP-Cl n=6. p-value: Paired t-test with Welch’s correction. Right: Cumulative βCtn4A-DD fluorescence intensity of PA crypts in NvocTMP-Cl versus Vehicle-treated mSIOs with local (precursorPA, budPA) 405nm irradiation (4s; 10 J/cm^2^). Boxplot 10-90%. Precursor PA: Vehicle n=8; NvocTMP-Cl n=8; bud PA: Vehicle n=6; NvocTMP-Cl n=8. p-value: Kruskal-Wallis, non-parametric ANOVA. **I.** Crypt growth as fold-change of crypt perimeter after 405 nm irradiation of NvocTMP-Cl (10 µM; blue, black) and Vehicle (light blue, grey) treated and locally 405 nm irradiated (10 J/cm^2^) bCtC-DD mSIOs. Mean+SD. NvocTMP-Cl : precursorPA n=8, budPA n=8; Vehicle: precursorPA n=8, budPA n=6.

**Fig. EV6.**
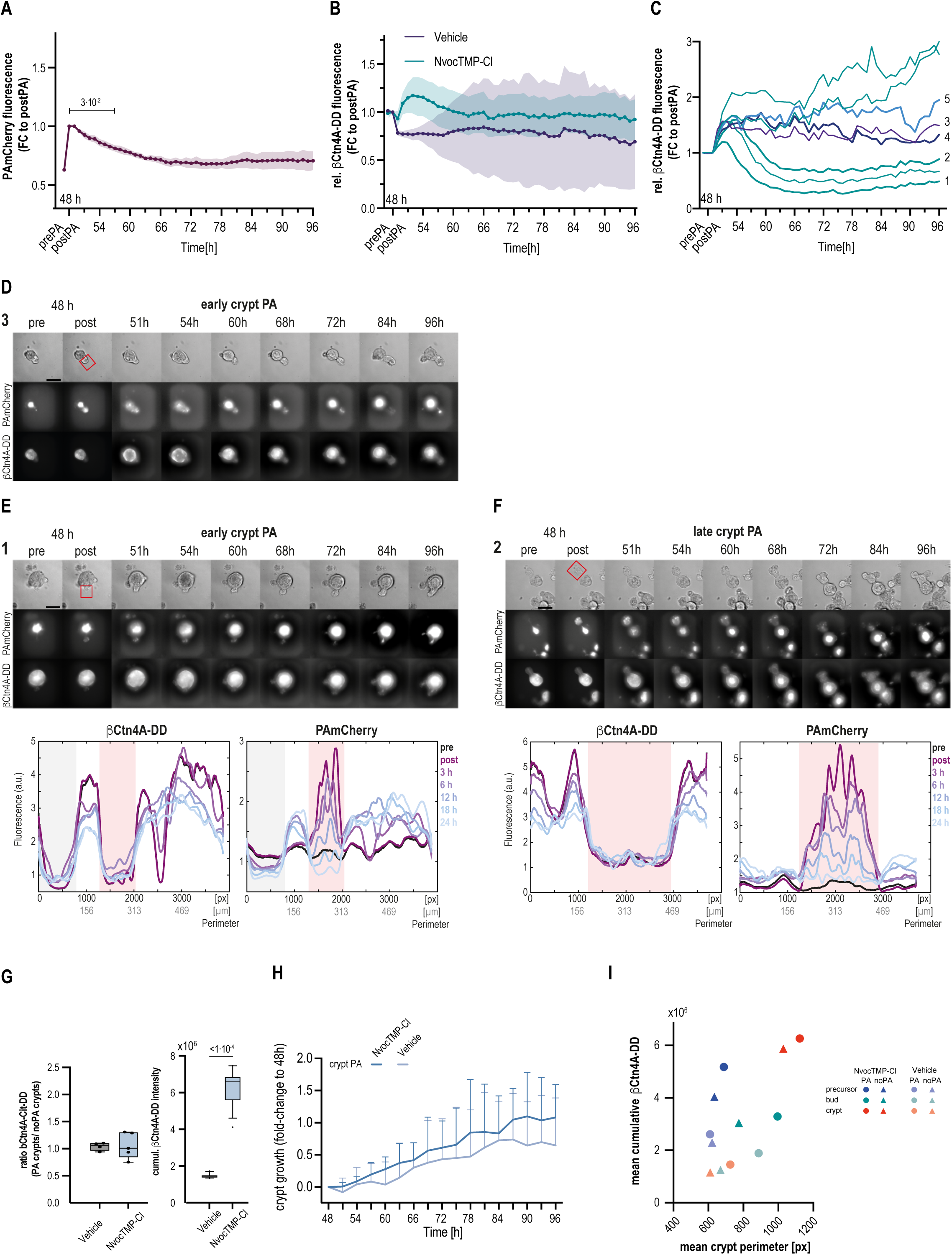
Disturbed organoid homeostasis by local opto-chemical oncoprotein activity and organoid homeostasis in controls. **A.** Expression of PAmCherry fluorescence over time normalized to postPA timepoint. Mean±95% CI. N=5; n=8. p-values: 2way ANOVA; Dunnett’s multiple comparison test compared to postPA. **B.** Relative expression of βCtn4A-DD in NvocTMP-Cl (10 µM) and Vehicle treated and locally 405 nm irradiated (4s; 10 J/cm^2^) bCtC-DD mSIOs over time, normalized to the postPA timepoint. Mean±95% CI. N=5/5; n=8/6. **C.** Single mSIO relative βCtn4A-DD expression ratio of NvocTMP-Cl (10 µM) treated to Vehicle-treated and locally 405 nm irradiated (4s; 10 J/cm^2^) bCtC-DD mSIOs over time, normalized to the postPA timepoint. Color corresponds to end-stage Mclass: blue: ‘Regular’, violet: ‘Crypty’, green: ‘CRYPT’, red: ‘Cystic’. Thick lines with numbers correspond to mSIO contours and MIP sequences in Fig. 6 D-H. **D.** Micrographs (maximum intensity projections) of exemplary developing bCtC-DD NvocTMP-Cl mSIO with local 405 nm irradiation in Transmission, PAmCherry and βCtn4A-DD pre and post local irradiation (PA; red rectangle) at indicated timepoints for early cryptPA (trajectory 3). **E.** Top: Micrographs (maximum intensity projections) of exemplary developing bCtC-DD Vehicle mSIO with local 405 nm irradiation in Transmission, PAmCherry and βCtn4A-DD pre and post local irradiation (PA; red rectangle) at indicated timepoints for early cryptPA (trajectory 1). Bottom: Linearized βCtn4A-DD and PAmCherry fluorescence along the developing mSIO contours for the NvocTMP-Cl treated examples 1 (early crypt PA) at indicated timepoints (prePA, postPA, 3 h, 6 h, 12 h, 18 h, 24 h) after PA with normalized and matched region sizes (villus and crypt regions at 18 h after PA). Background color indicates regions: red – PA region; grey – further crypts. **F.** Same as in A for the exemplary developing bCtC-DD Vehicle mSIO with late cryptPA (trajectory 2). **G.** Right: βCtn4A-DD fluorescence intensity ratio of PA versus non-PA crypts of the same mSIO. Boxplot Min-Max. cryptPA: Vehicle n=4; NvocTMP-Cl n=5. p-value: Paired t-test with Welch’s correction. Left: Cumulative βCtn4A-DD fluorescence intensity of PA crypts in NvocTMP-Cl versus Vehicle-treated mSIOs with local (cryptPA) 405nm irradiation (4s; 10 J/cm^2^). Boxplot 10-90%. cryptPA: Vehicle n=4; NvocTMP-Cl n=5. p-value: Kruskal-Wallis, non-parametric ANOVA. **H.** Crypt growth as fold-change of crypt perimeter after 405 nm irradiation of NvocTMP-Cl (10 µM; blue) and Vehicle (light blue) treated and locally 405 nm irradiated (10 J/cm^2^) bCtC-DD mSIOs. Mean+SD. NvocTMP-Cl : n=5; Vehicle: n=4. **I.** Scatterplot of mean cumulative βCtn4A-DD fluorescence versus mean crypt perimeter in Vehicle (light color) and NvocTMP-Cl-treated (darker color) and locally irradiated mSIOs crypt regions with PA (rectangle) versus noPA (triangle). Full scatter plots of all crypts over time in Supplements Fig. S6.

## Notes

### Competing Interest Statement

The authors have declared no competing interest.

### Summary of Updates

Correction of Colors in Figures Correction of In-text reference errors

## References

Amit S, Hatzubai A, Birman Y, Andersen JS, Ben-Shushan E, Mann M, Ben-Neriah Y, Alkalay I (2002) Axin-mediated CKI phosphorylation of beta-catenin at Ser 45: a molecular switch for the Wnt pathway. Genes Dev 16: 1066–1076

Andersson-Rolf A, Zilbauer M, Koo BK, Clevers H (2017) Stem Cells in Repair of Gastrointestinal Epithelia. Physiology (Bethesda) 32: 278–289

Andreu P, Colnot S, Godard C, Gad S, Chafey P, Niwa-Kawakita M, Laurent-Puig P, Kahn A, Robine S, Perret C et al (2005) Crypt-restricted proliferation and commitment to the Paneth cell lineage following Apc loss in the mouse intestine. Development 132: 1443–1451

Andreu P, Peignon G, Slomianny C, Taketo MM, Colnot S, Robine S, Lamarque D, Laurent-Puig P, Perret C, Romagnolo B (2008) A genetic study of the role of the Wnt/beta-catenin signalling in Paneth cell differentiation. Dev Biol 324: 288–296

Batlle E, Henderson JT, Beghtel H, van den Born MMW, Sancho E, Huls G, Meeldijk J, Robertson J, van de Wetering M, Pawson T et al (2002) β-catenin and TCF Mediate Cell Positioning in the Intestinal Epithelium by Controlling the Expression of EphB/EphrinB. Cell 111: 251–263

Becker WR, Nevins SA, Chen DC, Chiu R, Horning AM, Guha TK, Laquindanum R, Mills M, Chaib H, Ladabaum U et al (2022) Single-cell analyses define a continuum of cell state and composition changes in the malignant transformation of polyps to colorectal cancer. Nat Genet 54: 985–995

Behrens J, von Kries JP, Kühl M, Bruhn L, Wedlich D, Grosschedl R, Birchmeier W (1996) Functional interaction of beta-catenin with the transcription factor LEF-1. Nature 382: 638–642

Chen X, Venkatachalapathy M, Dehmelt L, Wu YW (2018) Multidirectional Activity Control of Cellular Processes by a Versatile Chemo-optogenetic Approach. Angew Chem Int Ed Engl 57: 11993–11997

Chen X, Venkatachalapathy M, Kamps D, Weigel S, Kumar R, Orlich M, Garrecht R, Hirtz M, Niemeyer CM, Wu YW et al (2017) "Molecular Activity Painting": Switch-like, Light-Controlled Perturbations inside Living Cells. Angew Chem Int Ed Engl 56: 5916–5920

Chen X, Wu YW (2018) Tunable and Photoswitchable Chemically Induced Dimerization for Chemo-optogenetic Control of Protein and Organelle Positioning. Angew Chem Int Ed Engl 57: 6796–6799

Clevers H (2026) Shifting paradigms in tissue stem cell biology: Insights from the intestine. Cell 189: 706–724

Clevers H, Nusse R (2012) Wnt/beta-catenin signaling and disease. Cell 149: 1192–1205

Clevers HC, Bevins CL (2013) Paneth cells: maestros of the small intestinal crypts. Annu Rev Physiol 75: 289-311

Date S, Sato T (2015) Mini-gut organoids: reconstitution of the stem cell niche. Annu Rev Cell Dev Biol 31: 269–289

Dawkins R (1976) The selfish gene. Oxford University Press, New York

De Landtsheer S, 2019. kmeans_opt, 1.0.0.1 ed., https://www.mathworks.com/matlabcentral/fileexchange/65823-kmeans_opt.

de Lau W, Barker N, Low TY, Koo BK, Li VS, Teunissen H, Kujala P, Haegebarth A, Peters PJ, van de Wetering M et al (2011) Lgr5 homologues associate with Wnt receptors and mediate R-spondin signalling. Nature 476: 293–297

Dekkers JF, Alieva M, Wellens LM, Ariese HCR, Jamieson PR, Vonk AM, Amatngalim GD, Hu H, Oost KC, Snippert HJG et al (2019) High-resolution 3D imaging of fixed and cleared organoids. Nat Protoc 14: 1756–1771

Donnelly MLL, Hughes LE, Luke G, Mendoza H, ten Dam E, Gani D, Ryan MD (2001a) The ’cleavage’ activities of foot-and-mouth disease virus 2A site-directed mutants and naturally occurring ’2A-like’ sequences. J Gen Virol 82: 1027-1041

Donnelly MLL, Luke G, Mehrotra A, Li XJ, Hughes LE, Gani D, Ryan MD (2001b) Analysis of the aphthovirus 2A/2B polyprotein ’cleavage’ mechanism indicates not a proteolytic reaction, but a novel translational effect: a putative ribosomal ’skip’. J Gen Virol 82: 1013–1025

Dow LE, O’Rourke KP, Simon J, Tschaharganeh DF, van Es JH, Clevers H, Lowe SW (2015) Apc Restoration Promotes Cellular Differentiation and Reestablishes Crypt Homeostasis in Colorectal Cancer. Cell 161: 1539–1552

Drost J, Clevers H (2018) Organoids in cancer research. Nat Rev Cancer 18: 407–418

Edelstein AD, Tsuchida MA, Amodaj N, Pinkard H, Vale RD, Stuurman N (2014) Advanced methods of microscope control using muManager software. J Biol Methods 1

Farrall AL, Riemer P, Leushacke M, Sreekumar A, Grimm C, Herrmann BG, Morkel M (2012) Wnt and BMP signals control intestinal adenoma cell fates. Int J Cancer 131: 2242–2252

Fearon ER (2011) Molecular genetics of colorectal cancer. Annu Rev Pathol 6: 479–507

Fearon ER, Vogelstein B (1990) A genetic model for colorectal tumorigenesis. Cell 61: 759-767

Fey SK, Vaquero-Siguero N, Jackstadt R (2024) Dark force rising: Reawakening and targeting of fetal-like stem cells in colorectal cancer. Cell Rep 43: 114270

Fujii M, Matano M, Nanki K, Sato T (2015) Efficient genetic engineering of human intestinal organoids using electroporation. Nat Protoc 10: 1474–1485

Harada N, Tamai Y, Ishikawa T, Sauer B, Takaku K, Oshima M, Taketo MM (1999) Intestinal polyposis in mice with a dominant stable mutation of the beta-catenin gene. Embo j 18: 5931–5942

Hartl L, Huelsz-Prince G, van Zon J, Tans SJ (2019) Apical constriction is necessary for crypt formation in small intestinal organoids. Dev Biol 450: 76–81

Hatzis P, van der Flier LG, van Driel MA, Guryev V, Nielsen F, Denissov S, Nijman IJ, Koster J, Santo EE, Welboren W et al (2008) Genome-Wide Pattern of TCF7L2/TCF4 Chromatin Occupancy in Colorectal Cancer Cells. Molecular and Cellular Biology 28: 2732–2744

Heinz MC, Oost KC, Snippert HJG (2020) Introducing the Stem Cell ASCL2 Reporter STAR into Intestinal Organoids. STAR Protoc 1: 100126

Hu M-K (1962) Visual pattern recognition by moment invariants. IRE Transactions on Information Theory 8: 179–187

Huang Z, Leng J, 2010. Analysis_of_Hus_moment_invariants_on_image_scaling_and_rotation, 2nd International Conference on Computer Engineering and Technology. Chengdu, China

Huber AH, Nelson WJ, Weis WI (1997) Three-dimensional structure of the armadillo repeat region of beta-catenin. Cell 90: 871–882

Huels DJ, Ridgway RA, Radulescu S, Leushacke M, Campbell AD, Biswas S, Leedham S, Serra S, Chetty R, Moreaux G et al (2015) E-cadherin can limit the transforming properties of activating beta-catenin mutations. EMBO J 34: 2321–2333

Huels DJ, Sansom OJ (2015) Stem vs non-stem cell origin of colorectal cancer. Br J Cancer 113: 1–5

Iwamoto M, Bjorklund T, Lundberg C, Kirik D, Wandless TJ (2010) A general chemical method to regulate protein stability in the mammalian central nervous system. Chem Biol 17: 981–988

Jacquemin G, Wurmser A, Huyghe M, Sun W, Homayed Z, Merle C, Perkins M, Qasrawi F, Richon S, Dingli F et al (2022) Paracrine signalling between intestinal epithelial and tumour cells induces a regenerative programme. Elife 11

Jarde T, Evans RJ, McQuillan KL, Parry L, Feng GJ, Alvares B, Clarke AR, Dale TC (2013) In vivo and in vitro models for the therapeutic targeting of Wnt signaling using a Tet-ODeltaN89beta-catenin system. Oncogene 32: 883–893

Kam Y, Quaranta V (2009) Cadherin-Bound β-catenin Feeds into the Wnt Pathway upon Adherens Junctions Dissociation: Evidence for an Intersection between β-catenin Pools. PLOS ONE 4: e4580

Kim S, Jeong S (2019) Mutation Hotspots in the β-catenin Gene: Lessons from the Human Cancer Genome Databases. Mol Cells 42: 8–16

Kim WK, Kwon Y, Jang M, Park M, Kim J, Cho S, Jang DG, Lee W-B, Jung SH, Choi HJ et al (2019) β-catenin activation down-regulates cell-cell junction-related genes and induces epithelial-to-mesenchymal transition in colorectal cancers. Scientific Reports 9: 18440

Langlands AJ, Almet AA, Appleton PL, Newton IP, Osborne JM, Nathke IS (2016) Paneth Cell-Rich Regions Separated by a Cluster of Lgr5+ Cells Initiate Crypt Fission in the Intestinal Stem Cell Niche. PLoS Biol 14: e1002491

Langlands AJ, Carroll TD, Chen Y, Nathke I (2018) Chir99021 and Valproic acid reduce the proliferative advantage of Apc mutant cells. Cell Death Dis 9: 255

Legland D, Arganda-Carreras I, Andrey P (2016) MorphoLibJ: integrated library and plugins for mathematical morphology with ImageJ. Bioinformatics 32: 3532–3534

Leng S, Pignatti E, Khetani RS, Shah MS, Xu S, Miao J, Taketo MM, Beuschlein F, Barrett PQ, Carlone DL et al (2020) beta-Catenin and FGFR2 regulate postnatal rosette-based adrenocortical morphogenesis. Nat Commun 11: 1680

Leslie A, Carey FA, Pratt NR, Steele RJ (2002) The colorectal adenoma-carcinoma sequence. Br J Surg 89: 845–860

Li VS, Ng SS, Boersema PJ, Low TY, Karthaus WR, Gerlach JP, Mohammed S, Heck AJ, Maurice MM, Mahmoudi T et al (2012) Wnt signaling through inhibition of beta-catenin degradation in an intact Axin1 complex. Cell 149: 1245–1256

Liu C, Li Y, Semenov M, Han C, Baeg G-H, Tan Y, Zhang Z, Lin X, He X (2002) Control of β-catenin Phosphorylation/Degradation by a Dual-Kinase Mechanism. Cell 108: 837–847

Liu Z, Chen O, Wall JBJ, Zheng M, Zhou Y, Wang L, Vaseghi HR, Qian L, Liu J (2017) Systematic comparison of 2A peptides for cloning multi-genes in a polycistronic vector. Sci Rep 7: 2193

MacDonald BT, Tamai K, He X (2009) Wnt/beta-catenin signaling: components, mechanisms, and diseases. Dev Cell 17: 9–26

Merker SR, Weitz J, Stange DE (2016) Gastrointestinal organoids: How they gut it out. Dev Biol 420: 239–250

Molenaar M, van de Wetering M, Oosterwegel M, Peterson-Maduro J, Godsave S, Korinek V, Roose J, Destrée O, Clevers H (1996) XTcf-3 transcription factor mediates beta-catenin-induced axis formation in Xenopus embryos. Cell 86: 391–399

Munemitsu S, Albert I, Souza B, Rubinfeld B, Polakis P (1995) Regulation of intracellular beta-catenin levels by the adenomatous polyposis coli (APC) tumor-suppressor protein. Proc Natl Acad Sci U S A 92: 3046–3050

Nusse YM, Savage AK, Marangoni P, Rosendahl-Huber AKM, Landman TA, de Sauvage FJ, Locksley RM, Klein OD (2018) Parasitic helminths induce fetal-like reversion in the intestinal stem cell niche. Nature 559: 109–113

Ozawa M, Baribault H, Kemler R (1989) The cytoplasmic domain of the cell adhesion molecule uvomorulin associates with three independent proteins structurally related in different species. Embo j 8: 1711–1717

Pan D, Klare K, Petrovic A, Take A, Walstein K, Singh P, Rondelet A, Bird AW, Musacchio A (2017) CDK-regulated dimerization of M18BP1 on a Mis18 hexamer is necessary for CENP-A loading. Elife 6

Pleguezuelos-Manzano C, Puschhof J, van den Brink S, Geurts V, Beumer J, Clevers H (2020) Establishment and Culture of Human Intestinal Organoids Derived from Adult Stem Cells. Curr Protoc Immunol 130: e106

Prost K, 2015. Development of a β-catenin stability sensor, Fakultät für Chemie und Chemische Biologie der Technischen Universität Dortmund. Technischen Universität Dortmund, p. 132.

Ray A (2021) Homeostasis and Homeorhesis: Sustaining Order and Normalcy in Human-Engineered Complex Systems. International Journal on Engineering Technologies and Informatics 2

Riemer P, Rydenfelt M, Marks M, van Eunen K, Thedieck K, Herrmann BG, Bluthgen N, Sers C, Morkel M (2017) Oncogenic beta-catenin and PIK3CA instruct network states and cancer phenotypes in intestinal organoids. J Cell Biol 216: 1567–1577

Saki F, Tahmasbi A, Soltanian-Zadeh H, Shokouhi SB (2013) Fast opposite weight learning rules with application in breast cancer diagnosis. Comput Biol Med 43: 32–41

Sansom OJ, Reed KR, Hayes AJ, Ireland H, Brinkmann H, Newton IP, Batlle E, Simon-Assmann P, Clevers H, Nathke IS et al (2004) Loss of Apc in vivo immediately perturbs Wnt signaling, differentiation, and migration. Genes Dev 18: 1385–1390

Sato T, Clevers H (2013) Growing self-organizing mini-guts from a single intestinal stem cell: mechanism and applications. Science 340: 1190–1194

Sato T, Stange DE, Ferrante M, Vries RG, Van Es JH, Van den Brink S, Van Houdt WJ, Pronk A, Van Gorp J, Siersema PD et al (2011a) Long-term expansion of epithelial organoids from human colon, adenoma, adenocarcinoma, and Barrett’s epithelium. Gastroenterology 141: 1762–1772

Sato T, van Es JH, Snippert HJ, Stange DE, Vries RG, van den Born M, Barker N, Shroyer NF, van de Wetering M, Clevers H (2011b) Paneth cells constitute the niche for Lgr5 stem cells in intestinal crypts. Nature 469: 415–418

Sato T, Vries RG, Snippert HJ, van de Wetering M, Barker N, Stange DE, van Es JH, Abo A, Kujala P, Peters PJ et al (2009) Single Lgr5 stem cells build crypt-villus structures in vitro without a mesenchymal niche. Nature 459: 262–265

Schindelin J, Arganda-Carreras I, Frise E, Kaynig V, Longair M, Pietzsch T, Preibisch S, Rueden C, Saalfeld S, Schmid B et al (2012) Fiji: an open-source platform for biological-image analysis. Nat Methods 9: 676-682

Serra D, Mayr U, Boni A, Lukonin I, Rempfler M, Challet Meylan L, Stadler MB, Strnad P, Papasaikas P, Vischi D et al (2019) Self-organization and symmetry breaking in intestinal organoid development. Nature 569: 66–72

Song JH, Huels DJ, Ridgway RA, Sansom OJ, Kholodenko BN, Kolch W, Cho KH (2014) The APC network regulates the removal of mutated cells from colonic crypts. Cell Rep 7: 94–103

Subach FV, Patterson GH, Manley S, Gillette JM, Lippincott-Schwartz J, Verkhusha VV (2009) Photoactivatable mCherry for high-resolution two-color fluorescence microscopy. Nat Methods 6: 153–159

Tahmasbi A, Saki F, Shokouhi SB (2011) Classification of benign and malignant masses based on Zernike moments. Comput Biol Med 41: 726–735

Tallapragada NP, Cambra HM, Wald T, Keough Jalbert S, Abraham DM, Klein OD, Klein AM (2021) Inflation-collapse dynamics drive patterning and morphogenesis in intestinal organoids. Cell Stem Cell 28: 1516–1532 e1514

Teague MR (1980) Image-Analysis Via the General-Theory of Moments. J Opt Soc Am 70: 920–930

Tsai CR, Galko MJ (2019) Casein kinase 1alpha decreases beta-catenin levels at adherens junctions to facilitate wound closure in Drosophila larvae. Development 146

Valenta T, Hausmann G, Basler K (2012) The many faces and functions of β-catenin . The EMBO Journal 31: 2714–2736

van der Wal T, van Amerongen R (2020) Walking the tight wire between cell adhesion and WNT signalling: a balancing act for beta-catenin. Open Biol 10: 200267

van Es JH, Jay P, Gregorieff A, van Gijn ME, Jonkheer S, Hatzis P, Thiele A, van den Born M, Begthel H, Brabletz T et al (2005) Wnt signalling induces maturation of Paneth cells in intestinal crypts. Nat Cell Biol 7: 381–386

Veeman MT, Slusarski DC, Kaykas A, Louie SH, Moon RT (2003) Zebrafish prickle, a modulator of noncanonical Wnt/Fz signaling, regulates gastrulation movements. Curr Biol 13: 680–685

Vermeulen L, Morrissey E, van der Heijden M, Nicholson AM, Sottoriva A, Buczacki S, Kemp R, Tavaré S, Winton DJ (2013) Defining stem cell dynamics in models of intestinal tumor initiation. Science 342: 995–998

Viragova S, Li D, Klein OD (2024) Activation of fetal-like molecular programs during regeneration in the intestine and beyond. Cell Stem Cell 31: 949–960

Waddington CH (2014) The strategy of the genes. Routledge

Wang BJ, Wang TT, Zhu HM, Yan R, Li XR, Zhang CQ, Tao WY, Ke XS, Hao PL, Qu Y (2022) Neddylation is essential for β-catenin degradation in Wnt signaling pathway. Cell Reports 38

Wang W, Lin C, Lu D, Ning Z, Cox T, Melvin D, Wang X, Bradley A, Liu P (2008) Chromosomal transposition of PiggyBac in mouse embryonic stem cells. Proc Natl Acad Sci U S A 105: 9290–9295

Wong SY, Chiam KH, Lim CT, Matsudaira P (2010) Computational model of cell positioning: directed and collective migration in the intestinal crypt epithelium. J R Soc Interface 7 Suppl 3: S351–363

Wu G, He X (2006) Threonine 41 in beta-catenin serves as a key phosphorylation relay residue in beta-catenin degradation. Biochemistry 45: 5319–5323

Xue SL, Yang Q, Liberali P, Hannezo E (2025) Mechanochemical bistability of intestinal organoids enables robust morphogenesis. Nat Phys 21: 608–617

Yang Q, Xue SL, Chan CJ, Rempfler M, Vischi D, Maurer-Gutierrez F, Hiiragi T, Hannezo E, Liberali P (2021) Cell fate coordinates mechano-osmotic forces in intestinal crypt formation. Nat Cell Biol 23: 733–744

Yui S, Azzolin L, Maimets M, Pedersen MT, Fordham RP, Hansen SL, Larsen HL, Guiu J, Alves MRP, Rundsten CF et al (2018) YAP/TAZ-Dependent Reprogramming of Colonic Epithelium Links ECM Remodeling to Tissue Regeneration. Cell Stem Cell 22: 35–49 e37

Zhu L, McNamara HM, Toettcher JE (2023) Light-switchable transcription factors obtained by direct screening in mammalian cells. Nat Commun 14: 3185

