## Appendix for "Oncomimetic β-catenin activity onset, duration and region defines aberrant intestinal development"

|| Authors contributed equally.
### deceased

Present address:
† Department of Molecular and Medical Virology, Ruhr University Bochum, Bochum, Germany
‡ Human BioMolecular Research Institute, San Diego, CA, USA
§ Institute for Physiology II, University of Bonn, Bonn, Germany

**Figure S5. Stages of mSIO development and visualization of βCtn4A-DD stabilization**

**A.** Micrographs of wildtype mSIO development over time from 24 h to 96 h. Developmental stages of developing crypts are indicated. Bulge (30 h): Region of columnar cells with Paneth cells which can be identified by granularity and color; Bud (36 h): Protrusion of the bulge region; Bud with apical constriction (48 h): clear apical constriction in the bud with lumen; Crypt (78 h): elongated bud with bottleneck forming at the hinge to the villus region. Villus states: fetal/regenerating 24/30 h lower right, 54/60 h upper left, characterized by thin cell layer; differentiated villus: thick columnar cell layer. Villus states switch during crypt formation and fission processes.

**B.** Developing mSIO contours colored by βCtn4A-DD fluorescence intensity for the NvocTMP-Cl precursorPA examples (Fig. 5 F,G; 1&2).

**C.** Developing mSIO contours colored by βCtn4A-DD fluorescence intensity for the NvocTMP-Cl budPA examples (Fig. 5 F,G; 3&4).

**D.** Developing mSIO contours colored by βCtn4A-DD fluorescence intensity for the Vehicle precursorPA and budPA examples (Fig. 5 E, Fig. EV5 F).

**Fig. S6. Scatterplots of cumulative βCtn4A-DD fluorescence versus crypt perimeter.**

**A.** Scatterplot of cumulative βCtn4A-DD fluorescence versus crypt perimeter for NvocTMP-Cl (circle) versus Vehicle (plus). Colors show individual bCtC-DD mSIOs over 48h after 405nm irradiation (4s; 10 J/cm$^2$). precursorPA (left), budPA (middle), cryptPA(right).

**B.** Scatterplot of cumulative βCtn4A-DD fluorescence versus crypt perimeter for NvocTMP-Cl PA crypts (circle) versus noPA crypts (plus). Colors show individual bCtC-DD mSIOs over 48h after 405nm irradiation (4s; 10 J/cm$^2$). precursorPA (left), budPA (middle), cryptPA(right).

**C.** Scatterplot of cumulative βCtn4A-DD fluorescence versus crypt perimeter for Vehicle PA crypts (circle) versus noPA crypts (plus).. Colors show individual bCtC-DD mSIOs over 48h after 405nm irradiation (4s; 10 J/cm$^2$). precursorPA (left), budPA (middle), cryptPA(right).

**Video1: bCtC organoid stably expressing bCtn4A-Cit in Transmission, 24-108h.**

**Video2: bCtC-DD organoid in Transmission, bCtn4A-Cit and PAmCherry after cryptPA, 48-99h.**

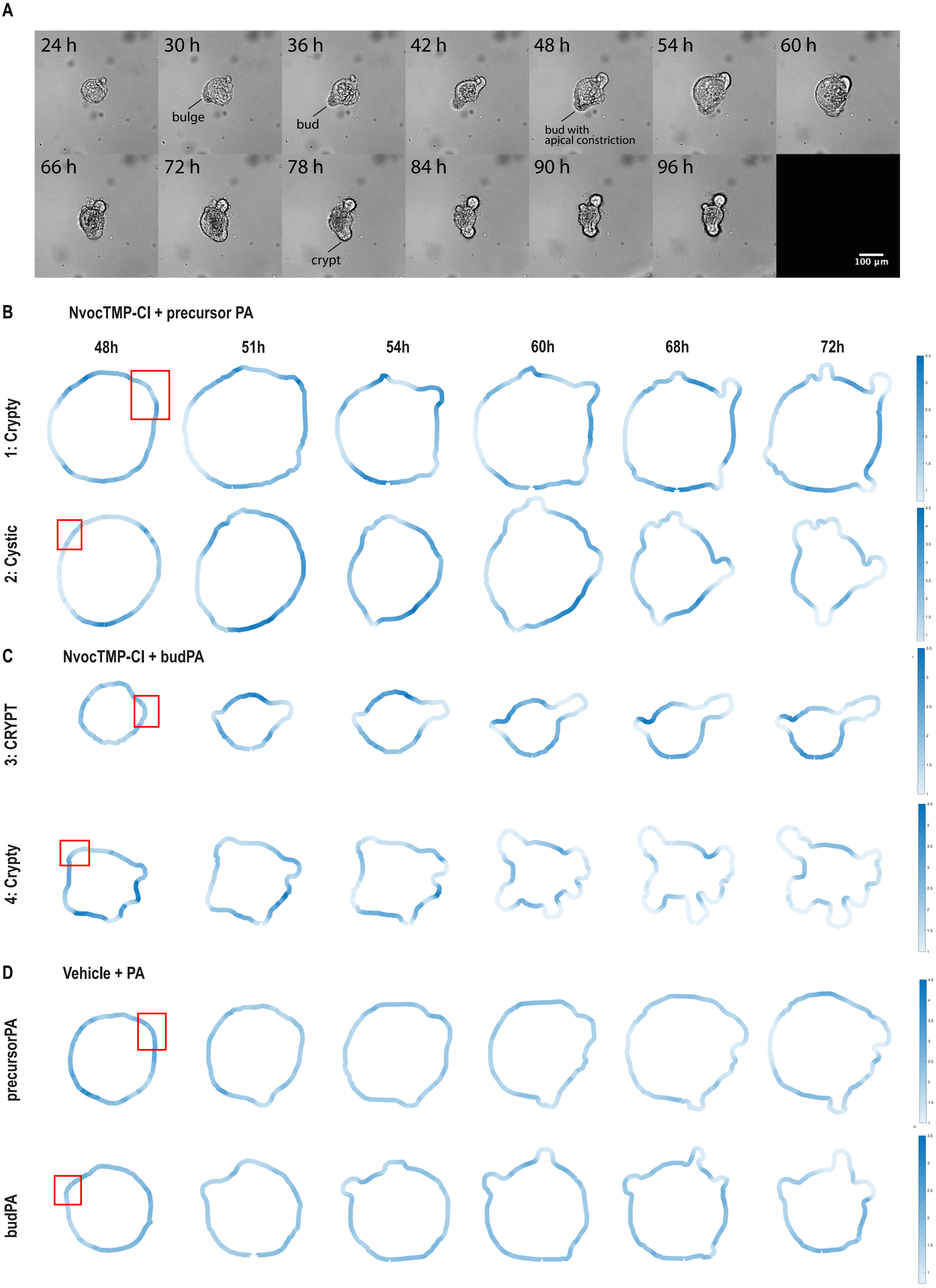
A
24 h
30 h
36 h
42 h
48 h
54 h
60 h
bulge
bud
bud with
apical constriction
66 h
72 h
78 h
84 h
90 h
96 h
crypt
100 µm
B
NvocTMP-Cl + precursor PA
48h
51h
54h
60h
68h
72h
1: Crypty
2: Cystic
C
NvocTMP-Cl + budPA
3: CRYPT
4: Crypty
D
Vehicle + PA
precursorPA
budPA

Figure S5

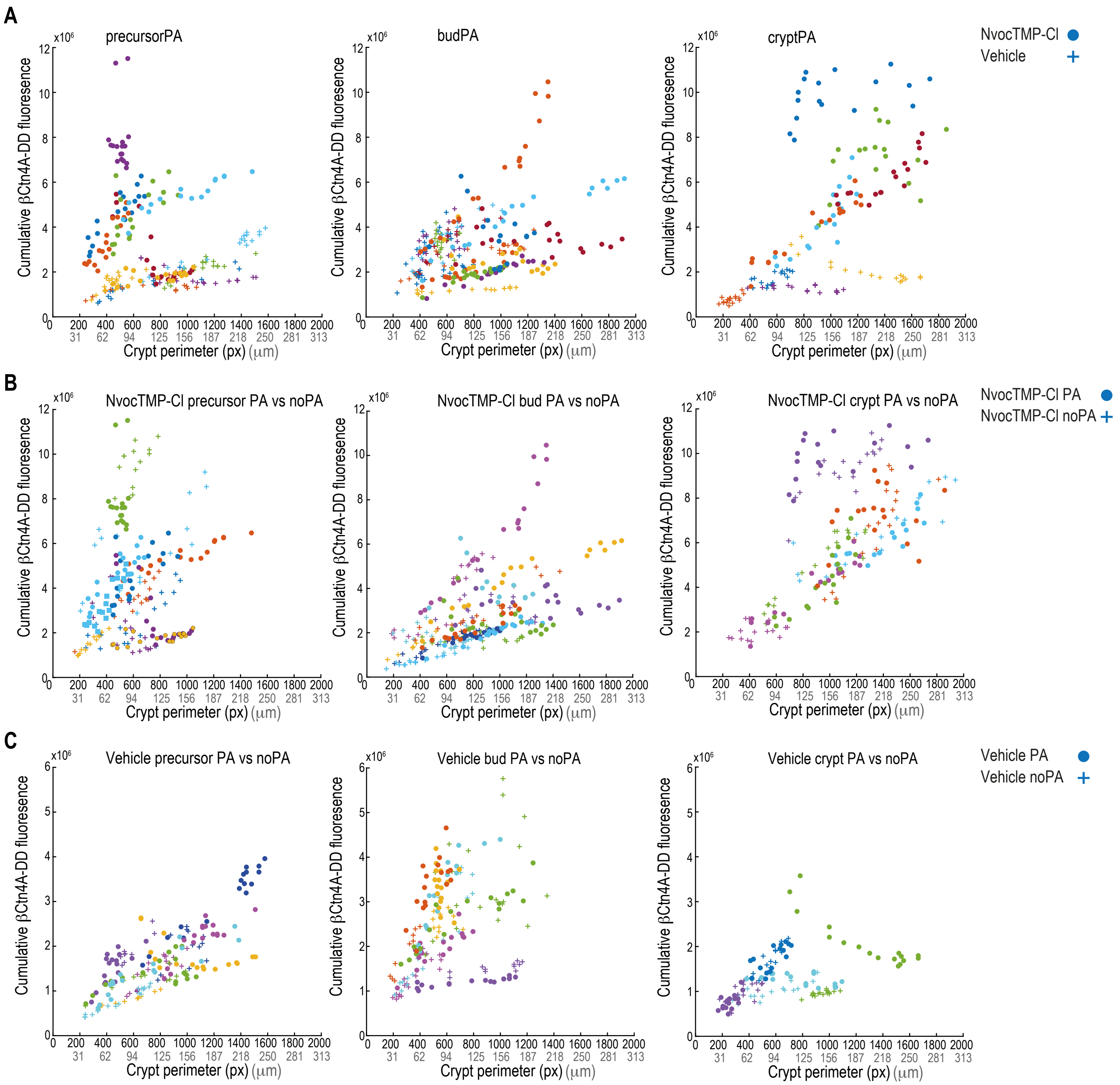
A
precursorPA
budPA
cryptPA
NvocTMP-Cl
Vehicle
B
NvocTMP-Cl precursor PA vs noPA
NvocTMP-Cl bud PA vs noPA
NvocTMP-Cl crypt PA vs noPA
NvocTMP-Cl PA
NvocTMP-Cl noPA
C
Vehicle precursor PA vs noPA
Vehicle bud PA vs noPA
Vehicle crypt PA vs noPA
Vehicle PA
Vehicle noPA
Cumulative βCtn4A-DD fluoresence
x10⁶
Crypt perimeter (px) (μm)

### Figure S6
